# VOLTAGE-GATED POTASSIUM CHANNELS CONTROL THE GAIN OF ROD-DRIVEN LIGHT RESPONSES IN MIXED-INPUT ON BIPOLAR CELLS

**DOI:** 10.1101/2020.10.10.334417

**Authors:** Christina Joselevitch, Jan Klooster, Maarten Kamermans

**Affiliations:** Department of Experimental Psychology, University of São Paulo, São Paulo, Brazil; Retinal Signal Processing, The Netherlands Institute for Neuroscience, Amsterdam, The Netherlands; Neurogenetics, Academic Medical Center, Amsterdam, The Netherlands

**Keywords:** scotopic vision, retina, bipolar cells, rods, potassium channels, gain control

## Abstract

To achieve high sensitivity at scotopic levels, vision sacrifices spatial and temporal resolution. The detection of dim light, however, depends crucially on the ability of the visual system to speed up rod signals as they advance towards the brain. At higher light levels, gain control mechanisms are necessary to prevent premature saturation of second-order neurons. We investigated how goldfish mixed-input ON bipolar cells (ON mBCs) manage to partially compensate for the intrinsically slow kinetics of rod signals in the dark-adapted state, and at the same time control the gain of rod signals. Rod-driven responses of axotomized ON mBCs become faster and more transient than those of rod horizontal cells as stimulus intensity increases. This transientness has a voltage-dependency consistent with the activation of a voltage-gated K^+^ conductance. Simulations with NEURON indicate that the voltage-gated K^+^ channels responsible for speeding up responses are concentrated at the distal tips of the bipolar cell dendrites, close to the glutamate receptors. These channels act as a gain control mechanism, by shunting the effect of tonically hyperpolarized rods onto the ON mBC. Further activation of K^+^ channels accelerates the ON mBC response by decreasing the membrane time constant as light levels increase. Therefore, the presence of voltage-gated K^+^ channels at the dendritic tips of ON mBCs extends the dynamic range of these neurons, and at the same time generates a transient signal already at the first visual synapse.

**Key Points Summary:**

- Here we show that voltage-gated potassium channels can adjust the gain of the rod input to mixed-input ON bipolar cells and generate a transient signal already at the first visual synapse.
- These channels are activated during the light-induced depolarization, making bipolar cell light responses smaller, faster, and more transient, effects that can be abolished by the K^+^ channel blocker TEA.
- Mathematical simulations suggest that these channels are concentrated at the bipolar cell dendritic tips, close to the site of rod input.
- This kind of gain control happens at all levels in the retina and is especially important for cells that receive mixed input from rods and cones, in order to prevent premature saturation with increasing light levels and remove the temporal redundancy of the photoreceptor signal.

## Introduction

The visual system needs to signal fast changes in local mean luminance and contrast to cope with the statistics of natural images and eye movements (Rieke and Rudd, 2009). At very dim light levels, vision is limited by fluctuations in photon absorption and quantal release, making integration over time and space a necessity (Balasubramanian and Sterling, 2009); as light levels increase, the temporal redundancy contained in sustained responses can be eliminated by making them transient. However, the makeup of neuronal networks could potentially slow down transmission of signals along the visual pathways. To compensate for delays generated by synaptic transmission and remove temporal redundancy from the neural code, retinal neurons at all layers have therefore several means of making their light responses transient.

Numerous such mechanisms were reported to exist in retinal bipolar cells. At the dendrites, depolarizing (ON) bipolar cells were shown to compensate for their intrinsically slow transduction cascade by a Ca^2+^-mediated feedback onto the light-driven conductance (Berntson et al., 2005; Shiells and Falk, 1999; Snellman and Nawy, 2002). Somatodendritic voltage-gated K^+^ channels were shown to decrease the time constant of the cell membrane (Mao et al., 1998; Mao et al., 2002), and in some species voltage-gated Na^+^ channels were also implicated in signal spread within subtypes of ON bipolar cells (Saszik and DeVries, 2012; Zenisek et al., 2001). Finally, mechanisms at the axon terminal, such as amacrine cell feedback (Kaneko and Tachibana, 1987; Shields et al., 2000), Ca^2+^ spikes (Dreosti et al., 2011; Protti et al., 2000; Zenisek and Matthews, 1997) and different types of feedback onto the voltage gated Ca^2+^ current (Palmer et al., 2003b; Palmer et al., 2003a), as well as intrinsic properties of the bipolar cell glutamate release (for examples, see Mennerick and Matthews, 1996 and Oesch and Diamond, 2011), would help make its output faster and more transient.

Despite the evidence for distal mechanisms that could potentially generate transientness (Hu and Pan, 2002; Klumpp et al., 1995; Mao et al., 1998; Mao et al., 2002; Pinto and Klumpp, 1998), rod-driven ON bipolar cell responses in some mammalian species are sustained (Euler and Masland, 2000; Oesch and Diamond, 2011). This would argue in favor of speed and transientness being generated at the bipolar cell output. Opposed to mammalian bipolar neurons, which are electrotonically compact (Oltedal *et al.*, 2009; but see Kuo et al., 2016), fish rod-driven bipolar cells are electrically coupled (Arai et al., 2010; Poznanski and Umino, 1997; Saito and Kujiraoka, 1988; Umino et al., 1994), implying considerable signal attenuation and decrease in response speed due to the large capacitance of the coupled network. In fish bipolar cells, therefore, additional mechanisms may be employed to speed up rod-driven signals before they reach the terminal.

We studied the rod-driven responses of mixed-input ON bipolar cells (ON mBCs) in the goldfish retina and found that dendritic voltage-gated K^+^ channels shape the kinetics of their light responses from high scotopic to mesopic light levels. These K^+^ currents speed up rod-driven light responses as light levels increase, reducing hereby temporal redundancy in the ON mBC output.

## Methods

### Ethical Approval

All animal experiments were carried out with permission of the ethical committee of the Royal Netherlands Academy of Arts and Sciences, acting in accordance with the European Communities Council Directive of 24 November 1986 (86/609/EEC).

### Retinal Slices

The procedure for making goldfish retinal slices has been described elsewhere (Joselevitch and Kamermans, 2007). Briefly, adult goldfish (*Carassius auratus*) in the light phase of their circadian rhythm were dark-adapted for 5-10 minutes prior to decapitation and enucleation under infrared illumination (λ > 850 nm). The anterior segment of the eye was removed, and the retina peeled off the pigment epithelium and sclera and placed receptor-side-up onto a piece of filter paper (13 mm diameter, 8 μm pore size, Millipore, Amsterdam, The Netherlands). Vacuum was applied to the other side of the filter in order to suck away the vitreous and attach the isolated retina firmly to the paper. The retina and the filter were cut in 100-150-μm-thick slices that were subsequently positioned on Vaseline tracks in a superfusion chamber for the electrophysiological experiments or processed for light/ electron microscopy.

### Solutions

The composition of the Ringer’s solution was (in mM): 102 NaCl, 2.6 KCl, 1 CaCl2, 1 MgCl2, 28 NaHCO3 and 5 glucose (248 mOsm). The solution was bubbled continuously with 2.5% CO_2_ and 97.5% O_2_ for a pH of 7.8. Picrotoxin (PTX, 100-500 µM) and strychnine (STRY, 5-18 µM) were added routinely to the control Ringer in order to block all GABAergic and glycinergic inputs to the recorded cells. For local drug delivery, DL-AP4 was dissolved in the control solution and pressure-applied locally with a puffer pipette.

The standard pipette solution contained (in mM): 0-60 KGlu or 0-94 KF, 10-50 KCl, 1 MgCl, 0.1 CaCl_2_, 10 ATP-Mg, 1 GTP-Na_3_, 0-10 cAMP, 0.1 cGMP-Na, 10 BAPTA-K_4_, 1 Lucifer Yellow-K_2_, 10-50 KOH (for pH 7.25, mOsm 212-273). No differences in cell behavior and/or response properties were observed with respect to these ranges of osmolarities and intracellular solution constituents, probably due to extensive coupling between bipolar cells in the dark-adapted retina (Arai et al., 2010). When needed, intracellular K^+^ channel block was attempted by replacing K^+^-based salts by equimolar Cs+ methanesulfonate and TEACl combinations in the pipette solution. Junction potentials for each pipette solution were calculated after Barry & Lynch (1991) and Ng & Barry (1995) and subtracted from the nominal voltage command values before correction for series resistance (see *Data Analysis* section). All chemicals were supplied by Sigma.

### Optical Stimulator

The superfusion chamber was mounted on a microscope equipped with infrared (λ > 800 nm) differential interference contrast optics (Eclipse E600-FN, NIKON, Japan) and the preparation was viewed in a TV monitor by means a 60 x water-immersion objective (N.A. 1.0) and a CCD camera (Philips, The Netherlands).

The light beam from one 400 W xenon arc and was separated into two optical paths with a beam splitter (Starna, UK). Both light beams passed through a series of neutral density filters (Schott, Germany), double interference filters (Melles Griot B.V., The Netherlands, peak transmission λs = 400, 450, 500, 550, 600, 650, 700 nm, 8 nm bandwidth), and circular variable neutral density filters (Barr & Stroud, UK).

Light stimuli were cast onto the retina via two different paths: small stimuli (slits of 50, 100 and 250 μm interposed on the optical path) were projected onto the preparation through the objective by means of mirrors and lenses, and a 3.5 cm field (referred to from now on as “full field”) was projected from below through the microscope condenser. Given the dimensions of the preparation, the effective diameter of the full field was equivalent to the width of the slices (3-6 mm). All set-up configurations were calibrated with a radiometer (50-245, irradiance head J1812, Tektronix, UK), and later with a photodiode (BPW34, Siemens, Germany). Cone light responses to stimulation originating from both channels were used as a control for the calibrations. Absolute intensity values are given in log quanta*μm^−^2*s^−1^.

### Electrodes and Recording Set-up

Patch pipettes were pulled from borosilicate capillaries (Harvard Apparatus Ltd., UK) with a Brown Flaming Puller (P-87, Sutter Instrument Co., Novato, CA) and had impedances between 5 and 15 MΩ when filled with pipette solution and measured in Ringer’s solution. Series resistances ranged from 6 to 33 MΩ and were corrected for junction potential and series resistance offline (see *Data Analysis* section). Electrodes were placed in a PCS-5000 micromanipulator (Burleigh Instruments, Inc., Fishers, NY) and connected to an Axopatch 200B Amplifier (Axon Instruments, Inc., Union City, CA). A second PCS-5000 micromanipulator was used to hold the puffer pipettes (manufactured as described above) and perfuse drugs locally by means of computer-controlled air pressure ejection.

Data acquisition and control of the optical stimulator were made by means of a Power 1401 AD/DA converter (Cambridge Electronic Design Ltd., UK) and an MS-DOS-based computer system or Signal 3 for Windows (Cambridge Electronic Design Ltd., UK). Voltage-clamp recordings were performed with corrected *V*_hold_ = −70 mV.

ON mBCs were visually selected based on their characteristic morphology and position in the outer part of the inner nuclear layer. Cell type was confirmed by measurement of response properties. Lucifer yellow was routinely included in the pipette solutions and dye-filled cells were observed immediately after the experiment. Data from both intact and axotomized cells were used. Unless otherwise stated, recordings from at least three cells yielded similar results for the experiments described.

### Data Analysis

Current transients were used to calculate series and input resistances. Current relaxations to −10-mV voltage steps from a holding potential of −70 mV were subtracted from the holding current and fit by double exponential functions of the form

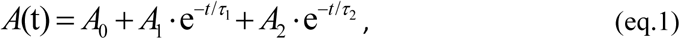

where *A*_0_ is the amplitude of the steady-state current at the end of the voltage step, *A*_1_ and *A*_2_ are the amplitudes of the two exponentials at the instant of voltage change, and *τ*_1_ and *τ*_2_ are the corresponding time constants of the current decay towards *A*_0_ (Mennerick et al., 1997; Nadeau and Lester, 2000). Fits begun 35-45 μs after the voltage change to avoid contamination by residual system filtering and extrapolated back to the onset of the voltage step in order to estimate the instantaneous current (*A*_0_+*A*_1_+*A*_2_). The calculated curves were subtracted from the data and the residual plots were used to control the quality of the fits (Ellis and Duggleby, 1978; Mennerick et al., 1997).

Series resistances were then calculated using the equation (Nadeau and Lester, 2000)

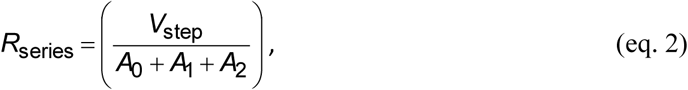

where *R*_series_ is the series resistance, and *V*_step_ is the applied voltage-clamp step. The voltage error for each recording was calculated offline according to the formula

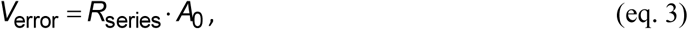

and subtracted from the nominal command voltages subsequently. Input resistances (*R*in) were estimated using the equation

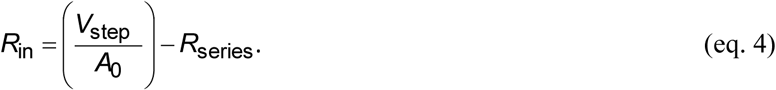

Voltage-gated currents were isolated by means of leak subtraction: the corrected whole-cell *IV* curves were subtracted from linear regressions of the current traces to voltage steps below –70 mV. Other forms of data analysis are mentioned in the text and figure legends.

### Anatomical Reconstruction

A retinal slice containing a Lucifer Yellow-filled ON mBC was fixed for 10 minutes at room temperature in 0.1 M phosphate-buffered (pH 6.5) 4% paraformaldehyde solution, washed in phosphate-buffered saline (PBS) for 10 minutes (2x), and blocked in 2% normal goat serum (NGS, Jackson ImmunoResearch Lab, West Grove, PA) in PBS for 20 min (Fig. 1A). The tissue was mounted on a poly-L-lysine coated slide (Menzel Gläser, Germany) and incubated overnight with a polyclonal antibody against Lucifer Yellow (anti-Lucifer Yellow, rabbit polyclonal, Millipore, 1:3000) at 4° C in PBS containing 0.3% Triton X-100 and 5% NGS. After washing (15 min, 3x, in PBS), the section was incubated in the secondary antibody (goat anti-rabbit Cy3, Jackson ImmunoResearch Lab, West Grove, PA; 1:500) for 30 minutes. The slide was cover-slipped with Vectashield (Vector Labs, Burlingame, CA) and observed on an inverted Zeiss Axiovert 100 M microscope equipped with the LSM 510 META laser scanning confocal module (Zeiss, Germany). Light micrographs were acquired as TIFF files directly from the microscope.

**Figure 1.**
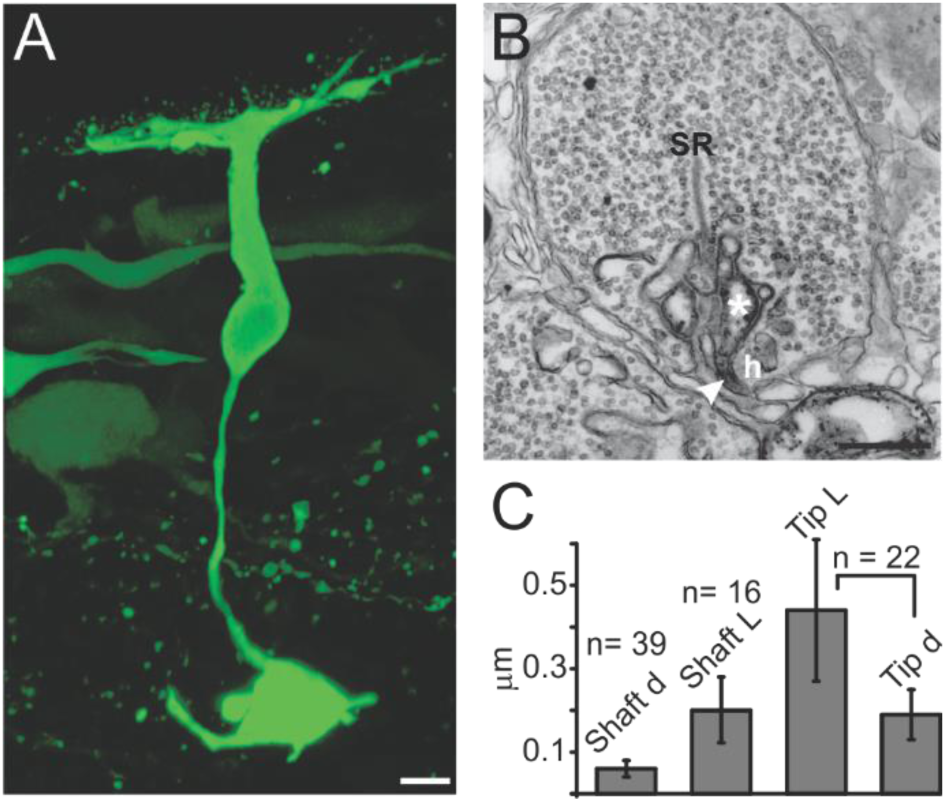
Anatomical reconstruction of a goldfish ON mBC. **A)** Lucifer-Yellow filled cell used for 3D reconstruction. Scale bar = 5 μm. **B)** EM picture of a rod terminal with PKC-immunoreactive processes used for estimating ON mBC dendritic and tip dimensions. Arrowhead = PKC-positive dendrite invaginating at the rod hilus (*h*); asterisk = dendritic tip of the same process opposite to a synaptic ribbon (*SR*). Scale bar = 500 nm. **C)** Mean values (± SD) for the diameter of PKC-positive processes at the rod hilus (*shaft d*, n = 39), the length of these processes (*shaft L*, n= 16), the diameter of PKC-positive dendritic tips opposite to synaptic ribbons (*tip d*, n = 22) and the lengths of the same tips (*tip L*, n = 22). All values are corrected assuming homogeneous shrinkage of 10% due to fixation and dehydration.

The TIFF files were then examined with the 3D-analysis software Image Pro (Media Cybernetics, Inc., Rockville, MD), and the diameters, lengths and 3D coordinates of the different compartments of the ON mBC were measured by means of a homemade analysis tool (NeuralDraw, written by Koos de Vos, *The Netherlands Institute for Neuroscience*). Because of the diminutive dimensions of the invaginating dendrites contacting rods, estimations of their diameters were made by means of measuring processes in 15 electronmicrographs of dendrites positively labeled for phosphokinase C (PKC), a marker for rod-driven ON bipolar cells (Fig. 1B and C). The starting values for the secondary dendrites and dendritic tips at cone terminals were obtained from a previous report (mb2s, Ishida *et al.*, 1980). The parameters obtained from this analysis and used in the mathematical model (described in the next section) are listed in Table 1.

**Table 1:** Dimensions of goldfish ON mBC compartments.

| Compartment | Length ($\mu\text{m}$) | Diameter ($\mu\text{m}$) | Source |
| --- | --- | --- | --- |
| Soma | 14 | 7 | **Fig. 1A** |
| Axon | 40 | 1 | **Fig. 1A** |
| Axon terminal | 7 | 7 | **Fig. 1A** |
| Primary dendrites | 26 | 1.3 | **Fig. 1A** |
| Secondary dendrites | 2 | 0.1* | Ishida *et al.* 1980** |
| Dendritic shaft | 0.2*** | 0.06* | EM pictures |
| Dendritic tip at rod terminals | 0.44 | 0.19 | EM pictures |
| Dendritic tip at cone terminals | 0.17 | 0.17 | Ishida *et al.* 1980** |

\* These values were varied in some simulations due to the uncertainty inherent to the morphological data.
\*\* Values correspond to those reported for type 2 ON mBCs (mb2s).
\*\*\* The length of the dendritic shaft was considered equivalent to the length of the rod hilus.

### Electron Microscopy

The same fixation was used as described for light microscopy. Transverse frozen sections, 30-40 µm thick, were obtained on a freezing microtome and collected in phosphate buffer (PB) at room temperature, which is equivalent to freeze-thawing. Retinal sections were incubated for 72 h with diluted antisera (anti-PKC, 1:400, mouse monoclonal, Sigma). After rinsing, the tissue was incubated in a Poly-HRP-goat anti-mouse IgG (PowerVision, ImmunoVision Technologies Co., Springdale, AR). To visualize the peroxidase, the material was incubated in a Tris-HCl diaminobenzidine (DAB, 0.05%) solution containing 0.03% H_2_O_2_. The DAB reaction product was subsequently intensified by the gold-substituted silver peroxidase method (Gorcs *et al.*, 1986).

Sections were rinsed in sodium cacodylate buffer 0.1 M (pH 7.4) and post-fixed for 20 min in 1% OsO4 supplemented with 1.5 % potassium ferricyanide in sodium cacodylate buffer. After rinsing in sodium cacodylate buffer, the material was dehydrated and embedded in epoxy resin. Ultrathin sections were observed in a Philips EM201 electron microscope (Philips, The Netherlands) and/or Technai (CM) 12 electron microscope (FEI, The Netherlands). Sections examined with the Philips electron microscope were counter-stained with uranyl acetate and lead citrate.

### Sampling and Production of Photomicrographs

At least 20 retinal sections obtained from a minimum of five animals were used for a given experiment. About five photomicrographs were taken from the most representative sections in each experiment. Light micrographs were acquired as TIFF files from the Zeiss LSM 510 META (1048×1024 pixels, 72 ppi). For EM experiments, the same number of retinal sections/animals was examined at the LM level; labeled sections were chosen for ultra-thin sectioning. A minimum of 40 photoreceptor terminals were observed per ultra-thin section. Ten to 30 micrographs were taken from these sections, depending on the presence and amount of labeling.

Electron micrographs obtained with the FEI electron microscope were directly acquired as TIFF files (1024×1024 pixels, 72 ppi). Electron micrographs obtained from the Philips electron microscope were first printed from the negatives for analysis. The negatives of the prints that were selected for publication were scanned on a sprint scan 4000 scanner (Polaroid, Breda, The Netherlands) and acquired as TIFF files at 600 ppi. All TIFF files were resampled at 400 ppi and subsequently resized and optimized for brightness and contrast using Photoshop (Adobe Systems).

### Computer Simulations

Modeling was carried out using NEURON 6.1.2 (Hines and Carnevale, 1997; Hines and Carnevale, 2000) running on Windows XP, Vista or 7. Integration time steps ranged from 0.00625 to 0.025 ms. The ON mBC compartmental model had 7 main parts, subdivided in a total of 495 sections (Fig. 2A): (i) the soma (divided in 11 segments), (ii) the axon (11 segments), (iii) axon terminal (one segment), (iv) six primary dendrites (9 segments each), (v) 162 secondary dendrites (9 segments each), (vi) the shafts of the dendritic tips (three segments each), and (vii) the tips of the secondary dendrites (one segment each). To test for spatial accuracy, we compared the kinetics of this model with those of a similar model in which the number of segments was multiplied by three at all sections, which would reduce spatial errors by a factor of nine (Carnevale and Hines, 2006); increasing the number of segments does not change the results described here.

**Figure 2.**
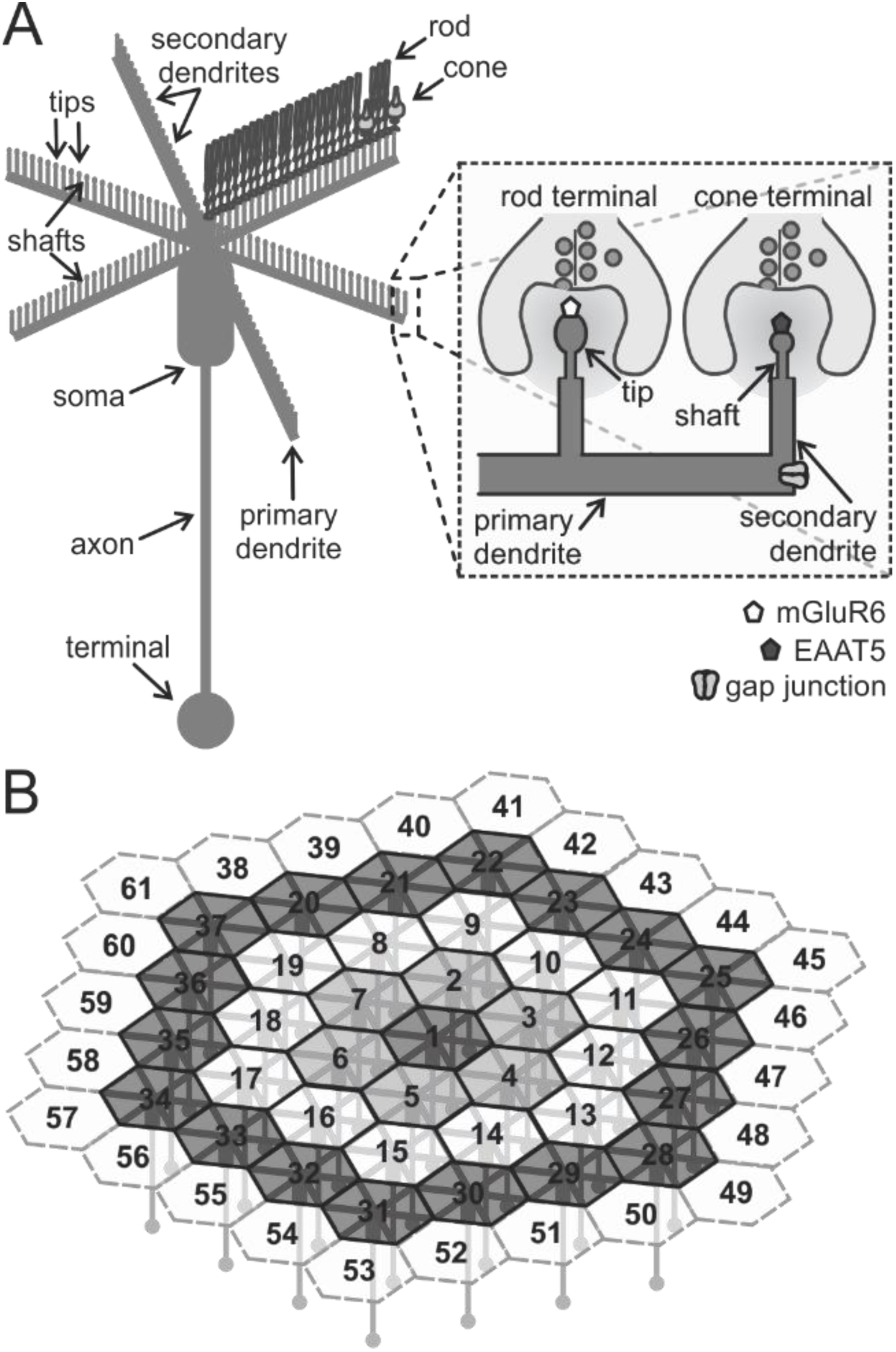
Schematic representation of the model ON mBC network. **A)** Schematic of the main compartments: axon terminal, axon, soma, primary dendrites, secondary dendrites, dendritic shafts, and tips (the three latter are not drawn to scale). Each primary dendrite contacts 25 rods and 2 cones (depicted in one of the primary dendrites as an example). The area within the dashed square is enlarged to show the position of the synaptic receptors at the tips of the dendrites and the extrasynaptic receptors at the secondary dendrites. Rods drive a conductance with *E*_rod_ = 0 mV (mGluR6), whereas cones drive a conductance with *E*_cone_ = −60 mV (EAAT5). Gap junctions between ON mBCs are at the extremities of the primary dendrites. **B)** Organization of the coupled syncytium. A central ON mBC (1) is connected to three rings of cells in a hexagonal lattice. The outer ring is connected to mBCs kept at *V*_rest_ (dashed hexagons).

Six primary (thick) dendrites sprouted from the soma of the ON mBC; each primary dendrite had 27 secondary (thin) dendrites attached to it: two contacting cones and 25 contacting rods, with a total of 12 cones and 150 rods projecting to the cell via the tips of the secondary dendrites. These numbers were chosen such as to yield a rod:cone ratio of 12.5, close to the value reported in literature for the goldfish mb2 (12.6, Ishida *et al.*, 1980). The total number of photoreceptors in the model (162) is however smaller than the one reported for intact cells (approximately 240), to account for loss of dendrites during the slicing procedure.

ON mBC membrane parameters, such as cytoplasmic resistivity (*R*_a_ = 250 Ω*cm, Mennerick *et al.*, 1997), input resistance (*R*_in_), specific membrane resistance (*R*_m_ = 67 kΩ*cm^2^) and specific membrane capacitance (*C*_m_ = 1μF/cm^2^) were assumed to be uniform throughout the cell, with a passive reversal potential of *E*_leak_ = −80 mV and a potassium equilibrium potential of *E*_K_ = −95 mV. Rods and cones, which provide synaptic input to the ON mBC, were modeled as 3-compartment cells (soma, axon and axon terminal, one segment each) with only passive conductances (*R*_a_ = 123 Ω*cm, *C*_m_ = 10 μF/cm^2^, *R*_m_ = 1000 Ω*cm^2^, *E*_leak_ = −50 mV).

Control of photoreceptor membrane potential was achieved by means of a simulated somatic voltage-clamp. Photoreceptor inputs were built such that rod hyperpolarization would drive a conductance increase in ON mBC with reversal potential *E*_rod_ = 0 mV, and cone hyperpolarization would drive a conductance decrease with *E*_cone_ = −60 mV (Grant and Dowling, 1995; Grant and Dowling, 1996). Synaptic conductances activated by photoreceptor polarization were simulated to yield light responses of physiological amplitudes and a resting membrane potential of around – 24 mV in the absence of K^+^ conductances: the rod-driven synaptic conductance density was −125 nA/cm^2^ in darkness and −300 nA/cm^2^ in light, and the cone-driven synaptic conductance density was kept at 300 nA/cm^2^.

In order to study the spatial dependence of the ON mBC responses, which are electrically coupled in the dark-adapted state (Poznanski and Umino, 1997; Yamada and Saito, 1997), the model cell was connected via the primary dendrite to a ring of 6 other neurons, and these were in turn connected to three concentric rows of cells, yielding a lattice of 61 cells (Fig. 2B). This number of coupled cells was chosen such that the receptive field of the central neuron was not disturbed by border effects: the ON mBCs in the outer ring (dashed hexagons in Fig. 2B) were kept at their resting membrane potential (*V*_rest_ = −43 mV), such that only the 37 innermost cells would effectively be polarized by current injection in the central neuron. The gap junctional current was simulated using the equation

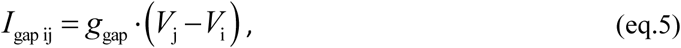

where *I*_gap ij_ is the gap junctional current flowing from the pre-synaptic cell to the post-synaptic cell, *g*_gap_ is the gap junctional conductance, *V*_i_ is the membrane potential of the post-synaptic cell, and *V*_j_ is the membrane potential of the pre-synaptic cell.

Voltage-gated K^+^ conductances (*I*_KV_) were simulated as Hodgkin-Huxley delayed rectifier (*m*^3*^*h*) currents (Fig. 3C, D and E). The location of K^+^ channels and their density were varied to reproduce the experimental data. Parameters were chosen such as to yield a voltage activation range and activation time constants similar to those recorded in real cells (Fig. 3A-B and E). However, due to the severe distortion of the *IV* relation and dynamic properties that remote conductances suffer when measured in non-spherical structures such as a bipolar cell (Hausser and Roth, 1997; Schaefer et al., 2003; Spruston et al., 1993), no further attempt was made to match the kinetics of the modeled currents to those measured in real cells.

**Figure 3.**
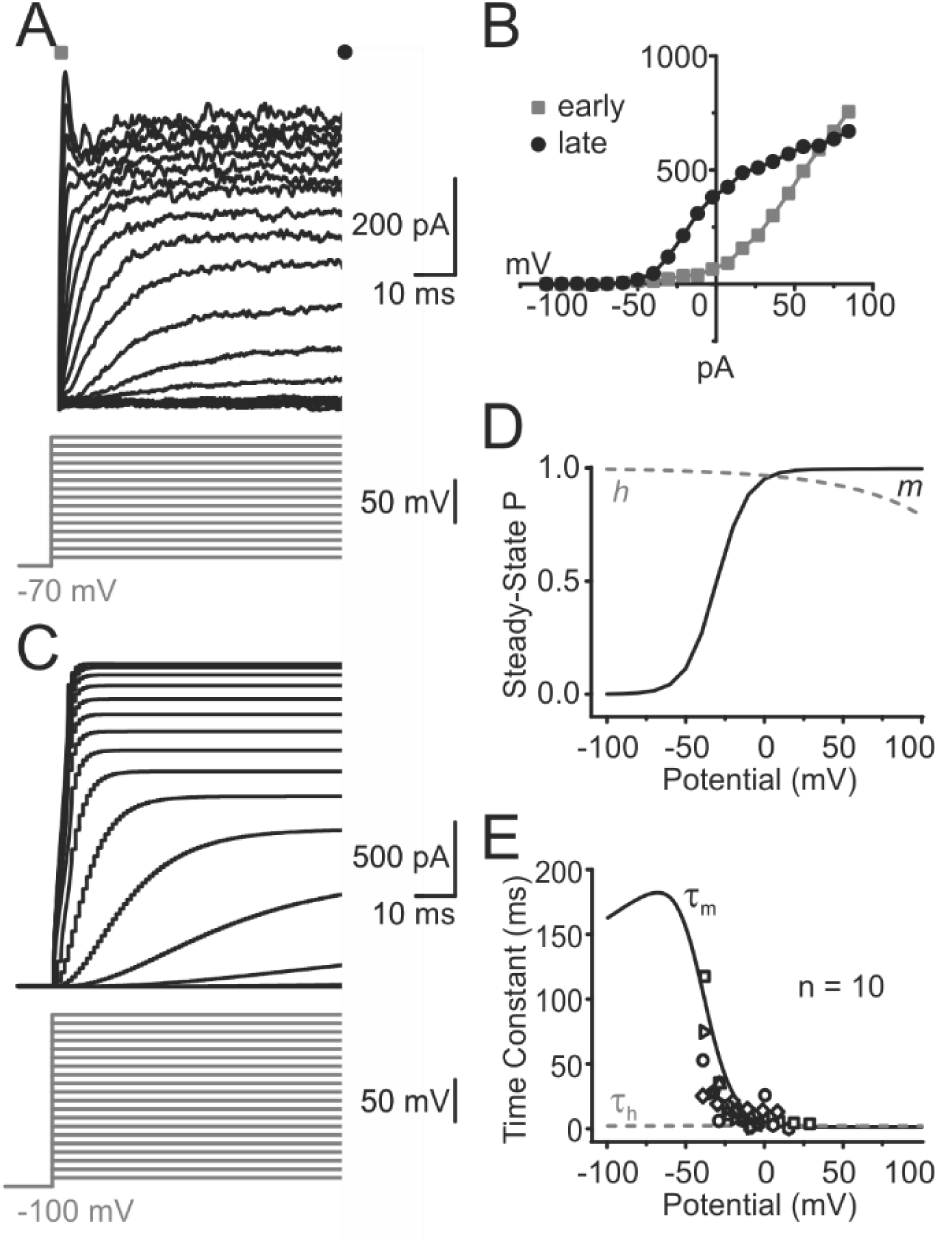
Native ON mBC voltage-gated K^+^ currents and the model *I*_KV_. **A)** Currents evoked in an ON mBC by 80-ms voltage steps from a holding potential of −70 mV, in 10 mV increments (black traces, top panel). The voltage-clamp protocol is shown in grey below the current traces. In this fast time scale, voltage-dependent outward currents have an early transient component (grey square) and a sustained component that inactivates little (black circle). **B)** Leak-subtracted *IV*-relations measured from the traces in (A) at the times indicated by the grey square and black circle. The transient component (grey squares) activates at potentials outside the physiological working range of the cell (−60 to −10 mV), whereas the sustained component (black circles) is active within this range. **C)** Responses of the model (black traces, top panel) to voltage steps from −90 to 100 mV from a holding potential of −100 mV (grey lines, bottom panel). The clamped structure is an isolated ON mBC axon terminal, with *I*_KV_ conductance density = 100 pS/μm^2^. **D)** Probability of steady-state activation (*m*, solid black line) and inactivation (*h*, dashed grey line) of the modeled currents. **E**, Time constant of activation (τ_m_, solid black line) and inactivation (τ_h_, dashed grey line). Open symbols are activation time constants obtained from exponential fits to voltage-clamp data of 10 goldfish ON mBCs measured in retinal slices.

### Statistics

Measured parameters were compared by means of one-way ANOVAs and bicaudal Student’s T-Tests with Bonferroni correction whenever applicable. Statistical significance was set at P ≤ 0.05.

## Results

### Characterization of ON mBC Rod-Driven Responses

Inputs from rods and cones are mediated by different mechanisms in fish ON mBCs: cone-driven light responses use glutamate transporters (EAATs, Grant and Dowling, 1996; Wong et al., 2005), whereas the rod-driven conductance is similar to that shown in the mammalian retina and is probably mediated by mGluR6 (Nawy and Copenhagen, 1987; Falk, 1988) and TRPM1 (Shen et al., 2009). To isolate the rod input to ON mBCs, we worked under dark-adapted conditions and used dim light stimuli that do not stimulate cones. In these circumstances, the spectral sensitivity of ON mBCs follows the rod absorption spectrum (Fig. 4A) and their absolute sensitivity is higher than that of cones (Joselevitch and Kamermans, 2007), indicating that their light responses are rod-driven.

**Figure 4.**
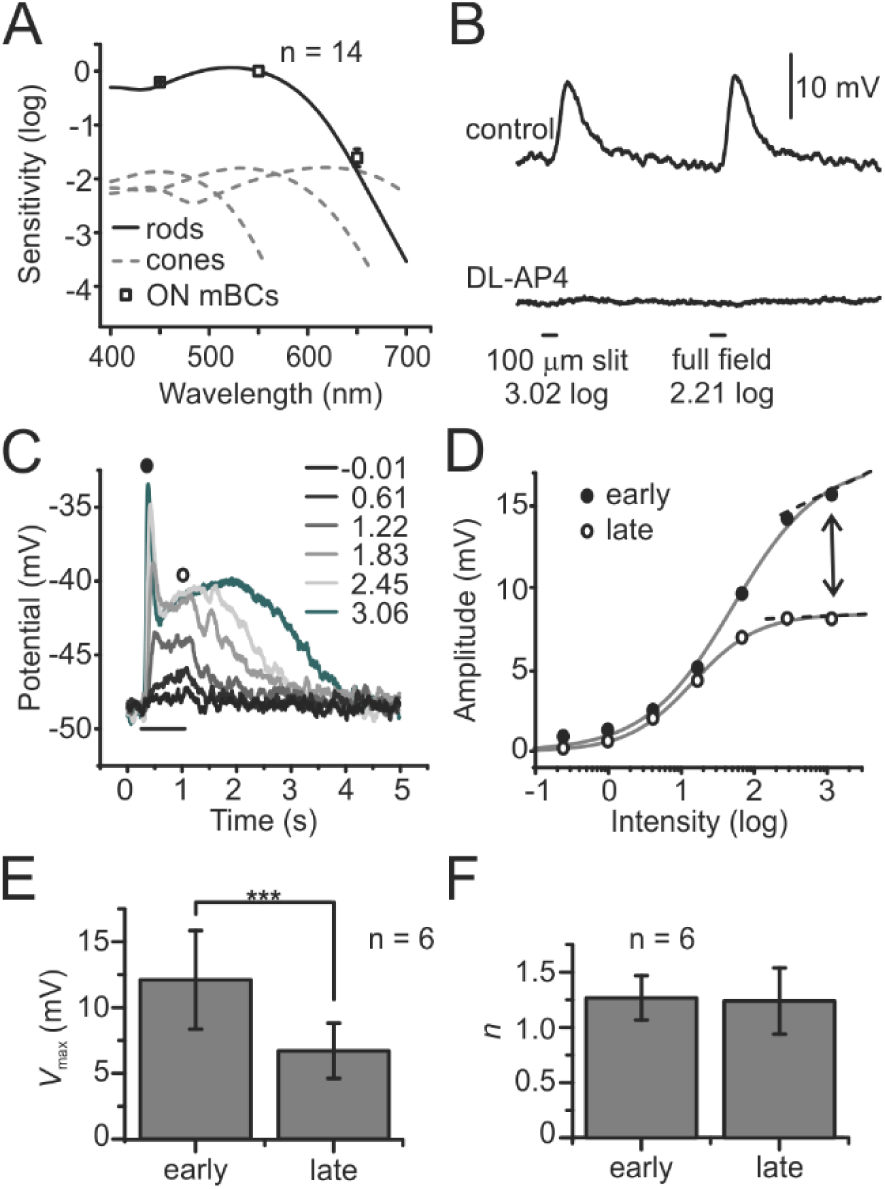
Characterization of rod-driven light responses in goldfish ON mBCs. **A)** ON mBCs are solely driven by rods in the dark-adapted retina. The mean (±SD) spectral sensitivity of dark-adapted ON mBCs follows the rod absorption spectrum (black line), and their absolute sensitivity is higher than that of S-, L- and M-cones (dashed grey lines). The stimulus was a 100 ms full-field light flash. ON mBC spectral sensitivity data was normalized to the rod peak sensitivity, cone spectra normalized to ON mBC sensitivity after measurements in 18 goldfish cones (Joselevitch and Kamermans, 2007). Photoreceptor spectra after Mooij & Van den Berg (1983). **B)** Rod-driven light responses are mediated by a group III mGluR. Bath application of 250 μM DL-AP4 hyperpolarized an ON mBC by 24 mV and completely suppressed the responses to 100 ms light flashes at 550 nm. Stimulus timing and intensity depicted in the figure. Similar results were obtained from 48 cells (DL-AP4, *n* = 44 cells; ACPT-1, *n* = 4). **C)** Rod-driven responses adapt during a light stimulus. Voltage responses of an axotomized ON mBC to 800 ms light stimuli of increasing intensities at 550 nm. Stimulus timing and intensity depicted in the figure. As intensity increases, light responses start displaying an initial transient peak (black circle) and a sustained plateau (open circle) at a more hyperpolarized level. **D)** Intensity-response relations of the cell in (C) measured at the times indicated by the black (early) and open (late) circles, respectively. At low intensities, there is little difference between the early and late components of the light response, whereas at higher intensities response amplitudes of the late component are smaller (arrow). Solid grey curves are Hill fits to the data points to obtain values for maximal response amplitudes (*V*_max_), sensitivity (*K*) and slope (*n*) of the curves. **E)** *V*_max_ (means ± SD) obtained in 6 axotomized cells (9 response families) for the same protocol as in (C). **F)** Slopes (means ± SD) obtained in 6 axotomized cells (9 response families) for the same protocol as in (C).

These rod-driven light responses are completely suppressed by group III mGluR agonists such as DL-AP4 or ACPT-1 (Fig. 4B) suggesting that they are solely mediated by mGluR6, as previously demonstrated in the mammalian retina (Slaughter and Miller, 1981; Yamashita and Wassle, 1991). Light responses of mammalian rod bipolar cells, however, were shown to be sustained (Euler and Masland, 2000; Oesch and Diamond, 2011, but see Berntson et al., 2005 and below), whereas those of ON mBCs become transient as intensity increases (Fig. 4C). To elucidate the functional consequences of this behavior, we measured the intensity-response relations early (150-250 ms) and late (400-500 ms) in the light response (solid and open circles in Fig. 4C-D, respectively), fitted Hill functions (grey curves in Fig. 4D) through these relations and determined their maximal response amplitudes (*V*_max_) and slope (*n*). The maximal amplitude (Fig. 4D, arrow, and Fig. 4E) was higher in the early phase compared to the late phase (P = 4*10^−5^), with no significant variation in slope (Fig. 4F). The change in response amplitude in time is equivalent to a temporal modulation of the rod-ON mBC synapse.

### Time- and Voltage-Dependent Changes in Rod-Driven Responses Are Due to ON mBC Intrinsic Properties

We subsequently investigated the mechanism underlying this transientness. To exclude a contribution of Ca^2+^ feedback onto the glutamate-gated channels (Berntson et al., 2005; Shiells and Falk, 1999; Snellman and Nawy, 2002), we added 10 mM of the Ca^2+^ buffer 1,2-bis(o-aminophenoxy)ethane-N,N,N’,N’-tetraacetic acid (BAPTA) to the patch pipette solution (Berntson *et al.*, 2005). Since under these conditions both intact (*n* = 13 cells, 20 response families) and axotomized cells (*n* = 13, 18 response families) still present transient light responses, we conclude that this transientness was unrelated to Ca^2+^-dependent processes or conductances located at the axon terminal. That left the somatodendritic compartment as a possible location, or the rod output itself. The latter option was studied next.

Although the rod-driven light responses of ON mBCs become transient with increasing intensity (Fig. 5A), this does not happen with responses of rod-driven horizontal cells (Fig. 5B). Since horizontal cells sense express AMPA-type glutamate receptors, which are much faster than the metabotropic receptors expressed in ON mBCs, these results suggest that the rod input to ON mBCs is sustained and does not become transient with increasing intensity. Therefore, the mechanism generating the transientness of rod-driven ON mBC responses must be intrinsic to bipolar cells.

**Figure 5.**
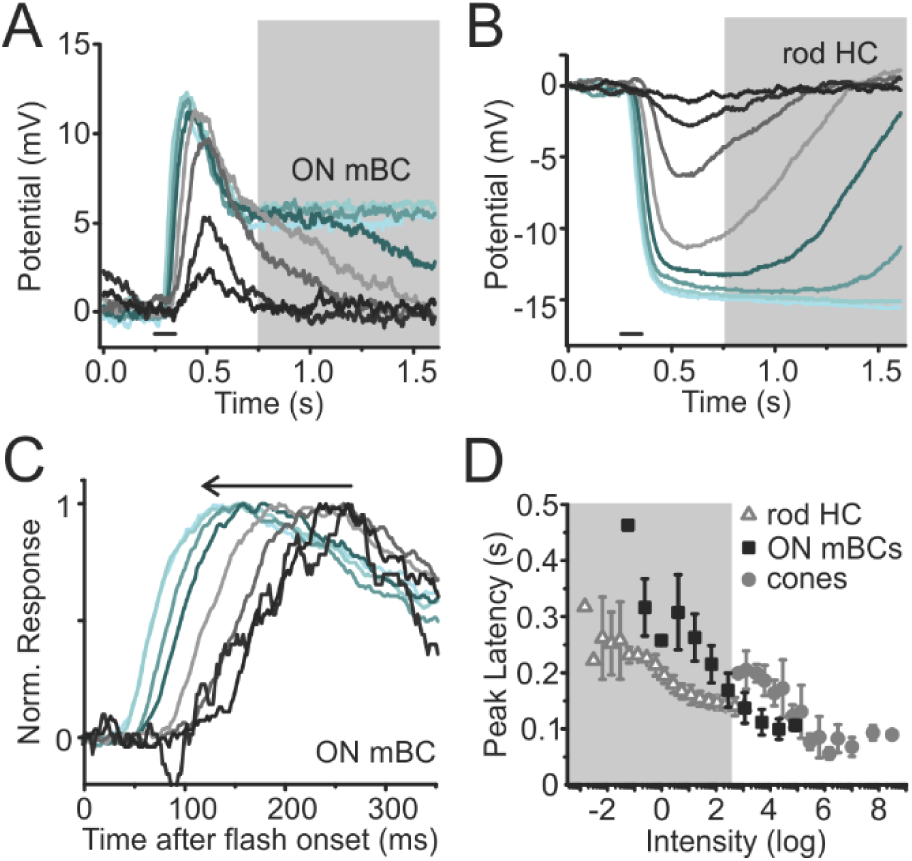
ON mBC light response kinetics derive from intrinsic properties. **A)** Responses of an ON mBC to 100 ms full-field light stimulation of increasing intensities at 550 nm (Intensities: 1.26, 1.92, 2.59, 3.26, 3.93, 4.57, 5.22 and 5.55 log quanta*μm^−2^*s^−1^). Stimulus timing depicted in the picture (black bar). As light intensity increases, a plateau with smaller amplitude develops after cessation of the light stimulus (shaded area). **B)** Rod-driven horizontal cell responses to the same stimuli as in (A) are sustained (shaded area), indicating that the output of goldfish rods to second-order neurons does not become transient with increasing light intensities. **C)** ON mBC responses become faster at higher light levels. Same responses of the cell in (A) in a faster time scale to show that as stimulus intensity increases, the cell reaches its maximal response sooner. **D)** Mean peak latency (± SD) for 6 ON mBCs (10 response families), 1 rod-driven HC (2 response families) and 7 cones (16 response families) for stimuli of increasing intensities at 550 nm. Peak latencies were measured from Boltzmann fits to the data; responses were considered to have reached their peak at 90 % of the curve maximum. The shaded area indicates intensities to which only rods are sensitive.

Not only do ON mBC responses become more transient with increasing light intensities, they also become substantially faster (Fig. 5C, arrow). Fig. 5D shows that this holds for ON mBCs, rod-driven horizontal cells and cones. To determine whether the reduction in time to peak originated in the photoreceptors, we plotted the peak latency for all three cell types as function of intensity and determined the slope of these curves, the temporal gain. The slopes were 62±14 ms/log quanta*μm^−2^*s^−1^ for ON mBCs, 64±18 ms/log quanta*μm^−2^*s^−1^ for cones and 38 ±5 ms/log quanta*μm^−2^*s^−1^ for rod-driven horizontal cells. Since both rod-driven horizontal cells and cones became faster at higher intensities, one can assume that part of this effect is a property of the photoreceptor response. The temporal gain of ON mBCs did not differ from that of cones, while it was higher than that of rod-driven HCs. Note however, that cones are only active at intensities in which ON mBCs are reaching saturation (non-shaded area in Fig. 5D and Fig. 6D). Conversely, at the intensity range in which cone and ON mBC responses overlap, ON mBCs are faster than cones (Fig. 5D). These findings again suggest that at least part of the ON mBC early response kinetics derive from their intrinsic properties and not primarily from their pre-synaptic inputs.

**Figure 6.**
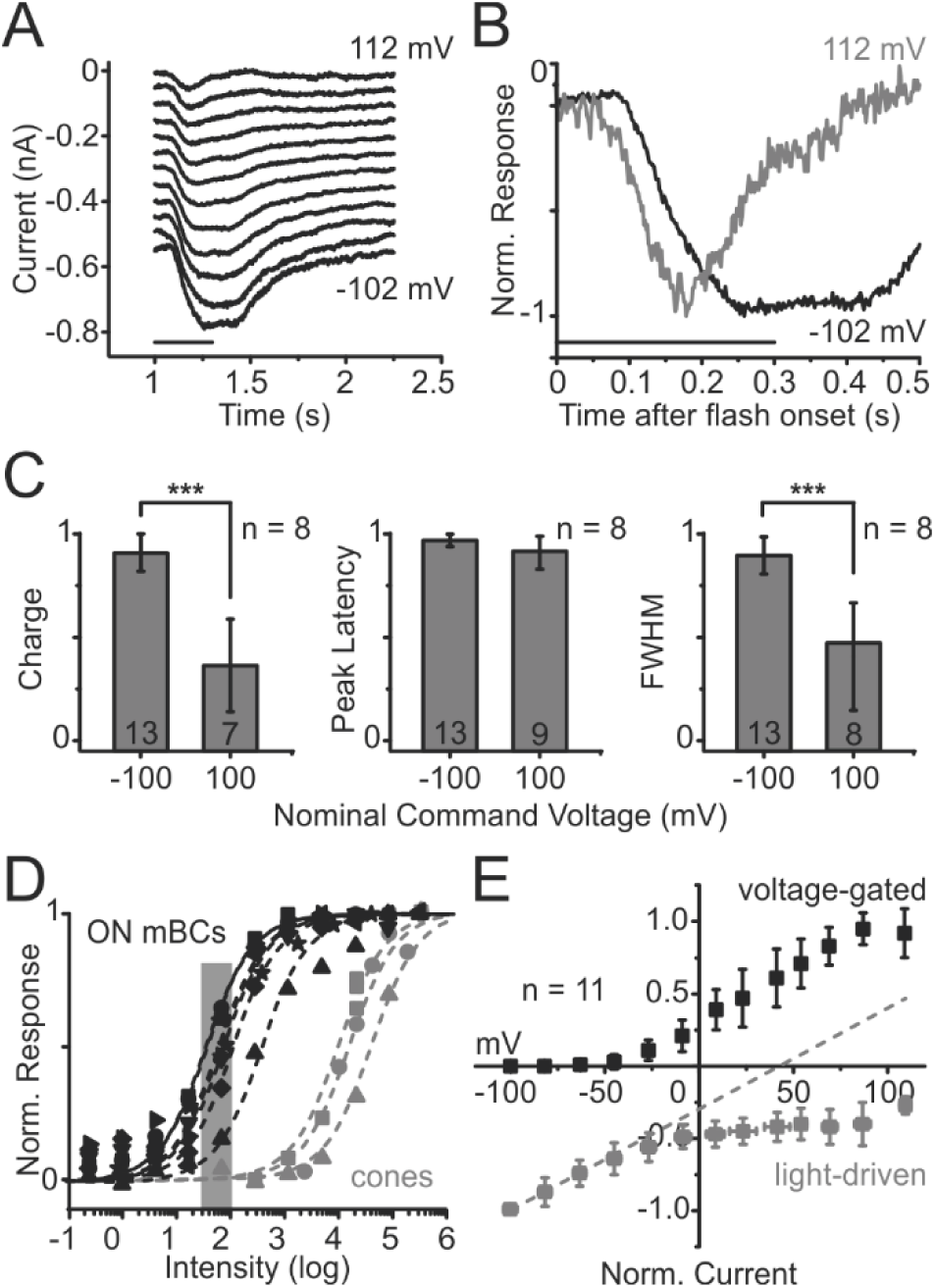
A voltage-gated K^+^ current interferes with the light-driven conductance. **A)** Unexpected absence of reversal for the light-driven conductance. Light-driven current responses of an axotomized ON mBC clamped at different potentials from a *V*_hold_ of −64 mV (from bottom to top trace: −102, −83, −63, −44, −25, −5, 14, 34, 53, 73, 92, 112 mV). Stimulus was a 1.83 log full-field flash at 550 nm. Light stimulus timing depicted in the figure. **B)** Potential-dependent kinetic changes of the light-driven current. The responses to −102 mV and 112 mV in (A) were normalized and depicted in a faster time scale to show that the light response is more transient at positive potentials. **C)** Summary of results obtained in 8 ON mBCs subjected to the protocol in (A). Current traces for nominal voltage steps to −100 mV and +100 mV from a nominal holding potential (*V*_hold_) of −60 mV (values not corrected for junction potential or series resistance, since these vary from cell to cell) were integrated to calculate the mean charge transfer ± SD (left panel), mean time-to-peak or peak latency ± SD (middle panel) and mean full width at half maximum ± SD (FWHM, right panel). The numbers inside each column in the graphs are the number of records used for each nominal command voltage. **D)** Absence of reversal potential for the light-driven conductance is not due to cone influence. Normalized intensity-response relations of the 8 ON mBCs in (C) and 3 cones (two M-cones and one L-cone) for full-field stimulation at 550 nm (black and grey symbols, respectively). Responses measured at the peak; dashed lines are Hill fits to the data. For most part of the dynamic range of ON mBCs, cones are still not active and cannot therefore account for the potential-dependent rectification and change in kinetics shown here. The shaded area highlights the stimulus intensities used in (A), (B) and (C). **E)** Rectification of light-driven *IV* relations in ON mBCs coincides with the activation of voltage-gated K^+^ currents. Mean normalized light-induced current (grey symbols) and mean normalized voltage-gated current (black symbols) of 11 axotomized ON mBCs (26 records). Light responses measured at the peak; voltage-gated currents calculated by subtracting the dark *IV*s from linear regressions extrapolated from data points more negative than the holding membrane potential (*V*_hold_). The dashed line shows that the light-induced slope conductance changes at around the same potential in which an outwardly rectifying voltage-gated current is activated.

An indication of the mechanism generating the transientness of the ON mBC response came from the *IV* relations of the light responses. Differently than expected based on literature (Berntson et al., 2004), light responses of goldfish ON mBCs did not reverse at 0 mV (Fig. 6A, *n = 81*). Instead, the *IV* relation of the light responses was inwardly rectifying, and the light response kinetics changed in a voltage dependent manner: they became more transient at positive potentials (Fig. 6B). Fig. 6C summarizes the results obtained in 8 ON mBCs. Responses at positive potentials are significantly smaller (P = 5.91*10^−6^) and more transient (P = 4.69*10^−6^), as indicated by the charge transfer and full width at half-maximal response (FWHM), respectively.

Because the pharmacology of these rod-driven responses is consistent with a single non-specific conductance and because cones are not responsive to the stimulus intensities used in these experiments (Fig. 6D), the lack of a clear reversal of the ON mBC light response is most likely due to poor space clamp of the dendrites (Spruston et al., 1993). Such space clamp problem could be caused by the activation of voltage-gated channels during depolarization, or due to gap-junctional coupling. We therefore studied first the effect of voltage-gated currents. The light-induced *IV* relations in ON mBCs (Fig. 6E, grey symbols) deviate from linearity at the same potential range in which outwardly rectifying channels are active (Fig. 6E, black symbols). We next set out to investigate whether these channels generate the intensity-dependent transientness of ON mBC light responses.

### K^+^ Channel Block Prevents Gain Control of Rod-Driven Light Responses

This outwardly rectifying voltage-gated current in bipolar cells has already been characterized as a delayed-rectifier, carried by K^+^ ions (Kaneko and Tachibana, 1985; Klumpp et al., 1995; Lasater, 1988; Tessier-Lavigne et al., 1988). To determine whether this kind of channel is essential for the kinetic changes in ON mBC responses, we blocked K^+^ channels by bath-applying the blocker tetraethylammonium (TEA, 10-50 mM). The choice of external block was made because use of Cs+ and TEA in the patch pipette failed to abolish the results shown so far, probably due to the fact that ON mBCs are electrically coupled (Arai et al., 2010) – signals recorded from one cell are contributed by the coupled network.

Fig. 7A-B shows that ON mBC light responses were larger and less transient when K^+^ channels were blocked (grey traces) compared to control (black traces). To quantify the effect of blocking K^+^ channels on the light responses, we integrated the light responses and obtained their area, peak latency and FWHM and compared the mean of three values pre-TEA and during TEA wash-in (Fig. 7C). Fig. 7D shows that in all cases, TEA increased the response amplitudes (> area, P = 2.55*10^−9^), slowed down responses (< peak latency, P = 2.65*10^−11^) and decreased transientness (> FWHM, P = 2.94*10^−7^). The resting membrane potential (*V*_rest_) of ON mBCs did not significantly change during TEA application (Fig. 7E). Although one would expect TEA to tonically depolarize ON mBCs, TEA also blocks K^+^ channels in photoreceptors (Bader et al., 1982; Fan and Yazulla, 1997), depolarizing them and thus leading to an increased basal glutamate release, which would in turn hyperpolarize ON mBCs. Together, TEA pre- and postsynaptic effects could lead to no net change in ON mBC resting membrane potential (Fig. 7E). The peak/plateau ratio increased significantly, indicating that the response became less transient (Fig. 7F). These results suggest that K^+^ channels are responsible for the transientness observed in rod-driven ON mBC light responses.

**Figure 7.**
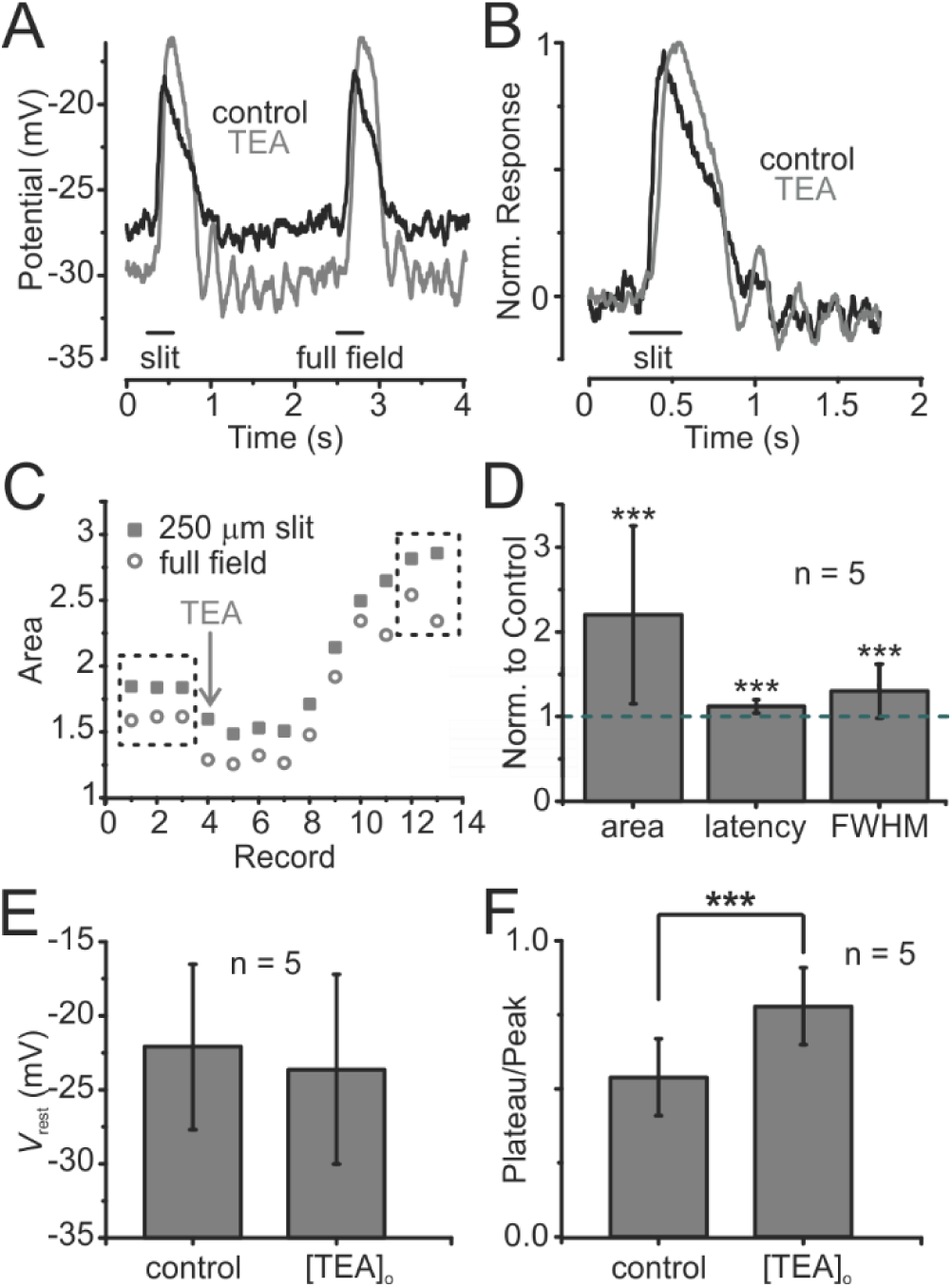
Voltage-gated K^+^ currents speed up ON mBC light responses. **A)** Responses of an ON mBC to two 250-ms light flashes at 550 nm (left: 250 μm slit, 2.58 log; right: full-field, 1.59 log) in control Ringer’s containing 250 μM PTX and 5 μM STRY to block GABAergic and glycinergic inputs (black trace) and in PTX+STRY+10 mM TEA to additionally block voltage-gated K^+^ currents (grey trace). Responses in TEA are larger, broader, and slower, suggesting that K^+^ currents make them faster and more transient, with little effect on resting membrane potential (*V*_rest_). **B)** Traces in (A) for stimulation with a slit are normalized and shown on a shorter time scale to emphasize that responses are slower and less transient in TEA. **C)** To quantify the effects of TEA, ON mBC responses before and during bath application of the drug were integrated and the area, peak latency and FWHM of the integrated responses were compared between the mean of three pre-TEA records and the mean of the last two records in TEA (dashed boxes) to determine how these parameters changed during wash in. **D)** Summary of results obtained in 5 cells normalized to pre-TEA values. Graphs show means ± SDs. **E)** Change in *V*_rest_ during the experiment was negligible in the same 5 neurons as in (D). **F)** Voltage-gated K^+^ channels make ON mBC light responses more transient. In external TEA, the relationship between response amplitudes measured at the plateaus (350-550 ms after light onset) and at the early peaks is significantly changed (P = 8.21*10^−4^): plateaus are about two times smaller in control (plateau/peak ratio = 0.54 ± 13 in control *versus* 0.78 ± 0.13, mean ± SD) and cells are more transient before TEA application.

### Voltage-Gated K^+^ Channels Are Concentrated in Distal Dendritic Compartments

The data so far suggest an intriguing interaction between the rod-driven glutamate-gated channels and K^+^ channels. To get a better understanding of the subcellular compartmentalization of these voltage-gated conductances and their possible interactions with the glutamatergic channels, we generated a model ON mBC. The influence of the physical localization of *I*_KV_ on the rod-driven *IV* relations of ON mBCs was investigated by means of five simulated conditions: (i) K^+^ channels localized only at the soma (Fig. 8A), (ii) K^+^ channels localized exclusively at the primary dendrites (Fig. 8B), (iii) K^+^ channels confined to the secondary dendrites (Fig. 8C), (iv) K^+^ channels localized only at the dendritic shafts (Fig. 8D), and (v) K^+^ channels localized exclusively at the tips of the secondary dendrites (Fig. 8E).

**Figure 8.**
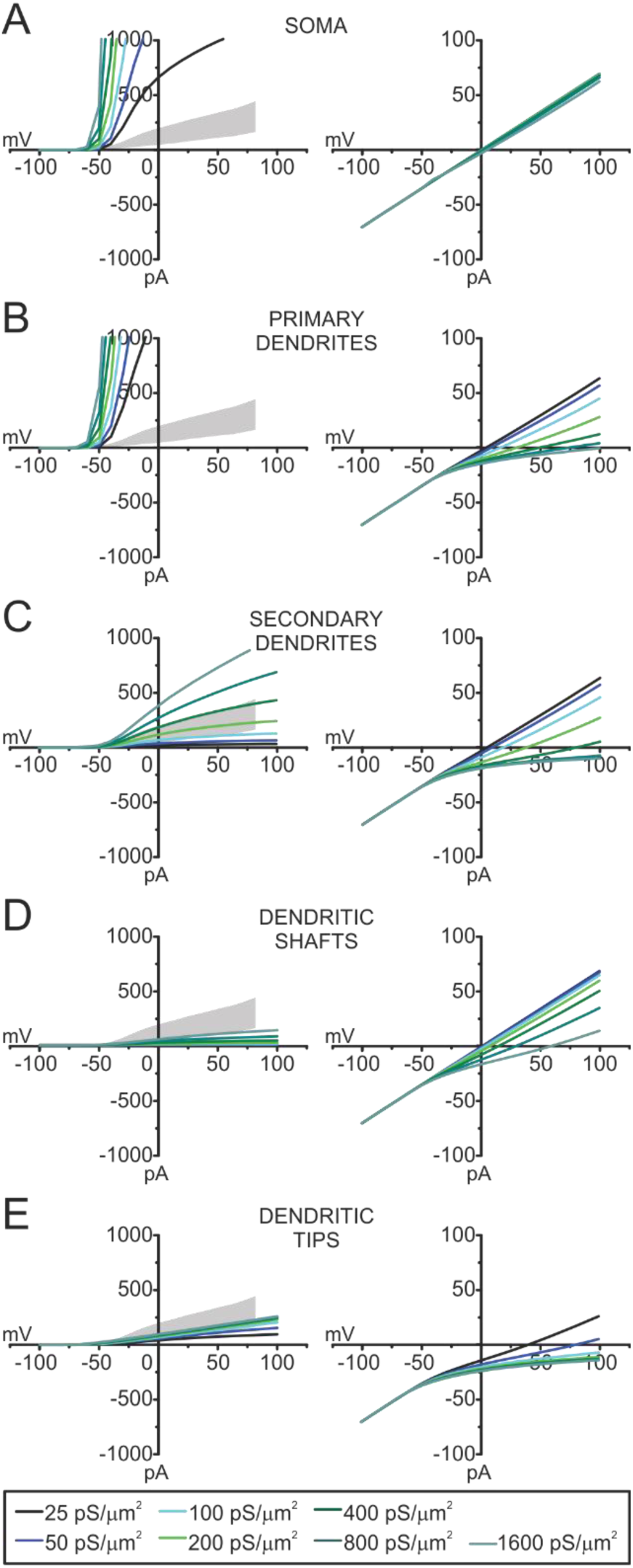
ON mBC voltage-gated K^+^ channels are concentrated in distal dendritic compartments. The graphs show the magnitude of voltage-gated currents (left) and their effect on the light-induced *IV*s (right) of a model ON mBC when *I*_KV_ is placed in different cell compartments. **A)** Soma. **B)** Primary dendrites. **C)** Secondary dendrites. **D)** Dendritic shafts. **E)** Dendritic tips. For all panels, *I*_KV_ conductance values are depicted at the bottom of the figure. Grey area represents mean ± SD voltage-gated currents of 28 records in 12 cells. For these simulations, the diameter of the secondary dendrites and dendritic shafts was set to 0.01 μm.

We then studied the relationship between the location and density of *I*_KV_, the non-linearity of the whole-cell *IV* relation and the rectification of the light response. When *I*_KV_ was located exclusively at the soma (Fig. 8A) or on the primary dendrites (Fig. 8B), voltage-gated currents (left panels) were always larger than those recorded in real cells (grey areas, *n* = 12). The *IV* relations of the light-driven conductance (right panels) were rather independent of *I*_KV_ when these channels were located in the soma or primary dendrites. Rectification was only present within the physiological range of voltage-gated *IV* relations when K^+^ channels were localized to more distal compartments such as the secondary dendrites (Fig. 8C), dendritic shafts (Fig. 8D) or to the tips of the dendrites (Fig. 8E). The largest rectification was observed when *I*_KV_ was confined to the tips of the dendrites (Fig. 8E), suggesting that a large part of these voltage-gated channels must be in close apposition to the sites of synaptic input.

### Dendritic Voltage-Gated K^+^ Channels Change the Kinetics of Rod-Driven Responses

We next investigated the effects of inserting *I*_KV_ at the dendritic tips on the kinetics of the rod-driven responses in current clamp and voltage-clamp modes. First, we gradually increased the polarization of the model rods, in order to simulate an intensity-response relation in the ON mBC in the absence and presence of a small K^+^ channel conductance at the dendritic tips (25 pS/μm^2^). In these simulations, dendritic *I*_KV_ increased the gain of the rod-ON mBC synapse (Fig. 9A): rod-driven ON mBC responses recorded at the soma became larger (grey traces) than when there was no *I*_KV_ (black traces), because of the *I*_KV_-induced change in *V*_rest_ (arrow) that increased the driving force for the rod-driven conductance.

**Figure 9.**
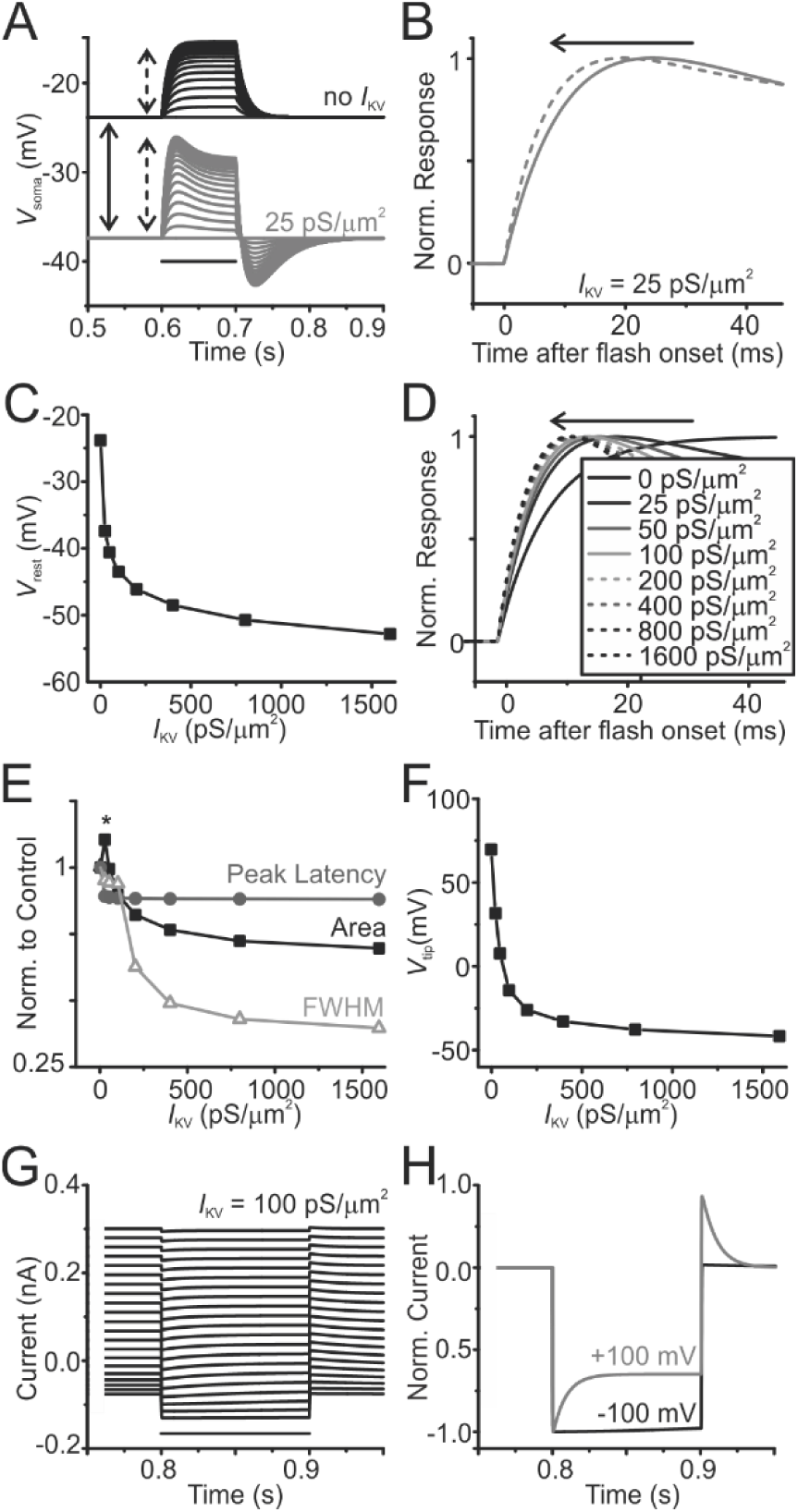
Dendritic *I*_KV_ changes the kinetics of ON mBC light responses. **A)** Somatic voltage responses (*V*_soma_) of the model ON mBC to a series of hyperpolarizing steps of the model rods when the cell lacks voltage-gated K^+^ channels (black traces) and with a 25 pS/μm2 voltage-gated K^+^ conductance inserted at the tips of the dendrites (grey traces). *I*_KV_ hyperpolarizes the model ON mBC (solid arrow) and makes light responses larger than when there is no *I*_KV_ (dashed arrows). **B)** Depolarization-induced activation of *I*_KV_ speeds up ON mBC responses. Normalized somatic voltage responses of the model ON mBC to the smallest and largest hyperpolarizing steps of the model rods shown in (A), with *I*_KV_ at the dendritic tips (conductance density depicted in the figure). Responses become faster as the light-driven depolarization increases (arrow). **C)** *V*_rest_ hyperpolarizes with increasing dendritic *I*_KV_ conductance density. **D)**y ON mBC response kinetics depend on *I*_KV_ density. Normalized somatic voltage responses of the model ON mBC to maximal hyperpolarization of the model rods, with increasing *I*_KV_ densities at the ON mBC dendritic tips (values in pS/μm2 depicted in figure). **E)** Changes in ON mBC response kinetics with increasing *I*_KV_ densities. Responses of the model ON mBC to maximal hyperpolarization of the model rods were integrated to investigate changes in response area (= amplitude), breadth (=FWHM) and speed (peak latency) elicited by changes in *I*_KV_ density at the dendritic tips in relation to control responses (no *I*_KV_). The asterisk denotes an initial increase in response amplitude due to the change in *V*_rest_. **F)** Dendritic *I*_KV_ prevents proper space clamp of remote compartments. Potential measured at the dendritic tip (*V*_tip_) of the model ON mBC during a somatic voltage step to +100 mV. **G)** *I*_KV_ is responsible for kinetic changes in light responses measured in voltage-clamp mode. Voltage-clamp experiment similar to the one in Fig. 6A. The model ON mBC was clamped from −60 mV to a series of voltage steps from −100 to +100 mV and stimulated with light (= hyperpolarization of the model rod) during the protocol. “Light” stimulus timing depicted as a horizontal bar in the picture. Similar to what happens in real cells, light responses do not reverse due to loss of space clamp and become more transient at positive potentials. *I*_KV_ conductance density depicted in the figure. **H)** Potential-dependent kinetic changes of the rod-driven current. The responses to −100 mV and 100 mV in (G) were normalized and depicted in a faster time scale to show that the light response of the model ON mBC is more transient at positive potentials. For these simulations, the diameter of the secondary dendrites and dendritic shafts was 0.01 μm.

This result might seemingly differ from the one presented in Fig. 7A-B, which show no net change in *V*_rest_ and an *increase* in ON mBC response amplitudes upon K^+^ channel block. However, one needs to take into consideration the fact that bath application of TEA has both pre- and post-synaptic effects, as explained previously, while our model includes only mechanisms intrinsic to the model bipolar cell. If the K^+^ channels are (partially) blocked intracellularly, ON mBCs indeed depolarize: cells recorded with Cs+- and/or TEA-based pipette solutions were about 20 mV more depolarized than those recorded with K^+^-based solutions (−26±16 mV *vs.* −45±16 mV, respectively, *n*1 =8, *n*2 = 13, P = 0.0168). Therefore, one expects that the presence of K^+^ channels in ON mBCs leads to hyperpolarization of these cells (Fig. 9A), with a consequent increase in the driving force for their glutamate-driven responses. Fig. 9B depicts the normalized minimal and maximal responses of the model cell for the grey traces in Fig. 9A, to illustrate that this small voltage-gated conductance density at the dendritic tips already speeds up rod-driven responses as the rod hyperpolarization increases.

We subsequently varied the dendritic *I*_KV_ conductance density to further investigate the effect of dendritic *I*_KV_ in the response kinetics of ON mBCs. Fig. 9C shows that increasing *I*_KV_ density at the dendritic tips indeed leads to hyperpolarization of the cell membrane. Furthermore, the response kinetics of the model ON mBC depended on *I*_KV_ density because time-to-peak for maximal rod hyperpolarization decreased as one increased the voltage-gated conductance density (Fig. 9D). These kinetic changes in current-clamp are summarized in Fig. 9E, which shows the effects of *I*_KV_ on response amplitude (area), speed (peak latency) and transientness (FWHM). Increasing dendritic *I*_KV_ density makes ON mBC light responses smaller (< area) and faster (< peak latency and FWHM).

We also tested whether this model could reproduce the voltage-dependent kinetic changes observed in voltage-clamp (Fig. 6). The potential at the dendritic tips (*V*_tip_) of the model ON mBC was investigated by applying a somatic voltage step from −60 to +100 mV and increasing the dendritic *I*_KV_ density (Fig. 9F). As expected, *I*_KV_ worsened the space clamp of the distal dendritic compartment. Finally, we simulated an experiment similar to that of Fig. 6A-B (Fig. 9G-H) and observed that this *I*_KV_-induced loss of space clamp indeed prevents the rod-driven responses of the model ON mBC from reversing at 0 mV, because the dendritic tips do not reach 0 mV. Furthermore, the responses of the model at more depolarized levels are more transient than the ones to hyperpolarizing steps (Fig. 9H), resembling our physiological observations. To summarize, our findings suggest that K^+^ channels are concentrated at the dendritic tips of ON mBCs and actively regulate the gain and speed of synaptic transmission from rods onto these neurons, by accelerating the time-to-peak of ON mBC light responses and their subsequent repolarization in an intensity-dependent manner.

### Electrical Coupling Increases Rectification

Since real ON mBCs are electrically coupled (Arai et al., 2010), and electrical coupling is known to induce space clamp problems such as the one reported here (Pang et al., 2004; Trexler et al., 2005), we proceeded to investigate the influence of electrical coupling in the light-induced currents of ON mBCs (see *Materials and Methods* and Fig. 2B). The single gap junctional range of conductances (*g*_gap_) used in the model are summarized in Fig. 10A, as well as the corresponding input resistance (*R*_in_) of the central cell for each *g*_gap_. These *g*_gap_ values were chosen such as to yield a receptive field diameter of approximately 370 µm, which is close to the lower values reported in literature for carp ON mBCs (from 300 µm to 1 mm; Saito and Kujiraoka, 1988), and for salamander ON BCs (from 345 to 662 µm; Borges and Wilson, 1987).

**Figure 10.**
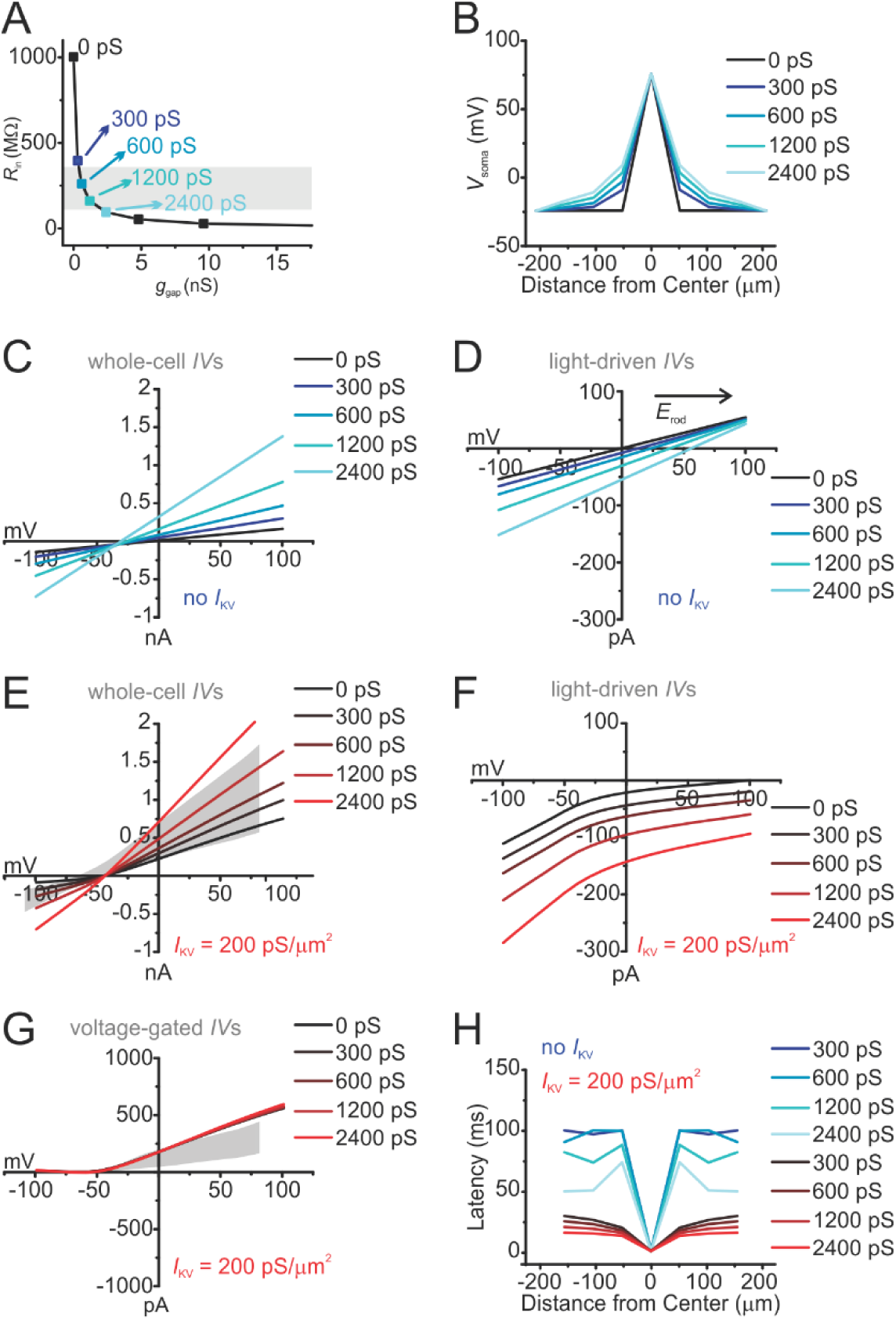
The influence of electrical coupling on the physiology of ON mBCs. **A)** Changes in input resistance (*R*_in_) with increased gap junctional coupling. The grey area shows mean ± SD *R*_in_ measured in 30 records from 8 axotomized ON mBCs. **B)** Electrical coupling increases the size of the ON mBC receptive field. Somatic membrane potentials of cells in different rings of the coupled syncytium during a +100 mV voltage step applied to the central ON mBC from *V*_rest_ (−24 mV, due to the absence of K^+^ conductances in this simulation). The amount of polarization of the neighboring cells increases with the strength of coupling (values of *g*_gap_ are depicted in the legend). **C)** Electrical coupling increases whole-cell currents. Whole-cell *IV* relations of the central ON mBC when clamped at a *V*hold of −70 mV and stepped from −100 to +100 mV for different strengths of coupling (values of *g*_gap_ are depicted at the right corner of this row). As the strength of coupling increases, *R*_in_ decreases and as a result whole-cell currents are augmented. **D)** Electrical coupling shifts the apparent reversal potential for the rod-driven currents (*E*_rod_). Light-induced *IV* curves measured in the central ON mBC for the simulation in (C). *E*_rod_ shifts proportionally towards more positive values as coupling is increased, but no rectification is induced. **E-G)** Same simulations as in (C-D), but with *I*_KV_ at the dendritic tips (200 pS/μm^2^). For these simulations, diameter of secondary dendrites = 0.02 μm, and diameter of the dendritic shafts = μm. Values of ggap are depicted in the legend. Grey area represents mean ± SD of 28 records in 12 cells. **F)** Electrical coupling increases the rectification induced by *I*_KV_. Simulations of the changes in the rod-driven *IV* relation of an ON mBC for full-field stimulation induced by varying the gap junctional conductance when K^+^ channels are placed at the dendritic tips. **G)** Voltage-gated *IV* relations for the simulation in (E). Grey area represents mean ± SD of 28 records in 12 cells. **H)** *I*_KV_ counteracts the slowing down in response kinetics caused by electrical coupling. Peak latency of light responses of ON mBCs in the coupled network for the simulation in (B) without dendritic *I*_KV_ (blue traces) and with *I*_KV_ at the dendritic tips (red traces). Ggap values depicted in the figure.

The spatial integration in the ON mBC network is illustrated in Fig. 10B, in which the somatic membrane potential (*V*_soma_) of cells in the different rings of the ON mBC lattice are depicted for a voltage step (*V*_step_) of 100 mV applied to the soma of the central cell. Neighboring cells are polarized to a smaller extent by the *V*_step_, and the amount of polarization is proportional to *g*_gap_. Consequently, as shown in Fig. 10C, electrical coupling leads to an increase in the amplitude of the ON mBC whole-cell currents. The reversal potential of all whole-cell currents is around −40 mV, because this is the resting membrane potential of the cells in the ON mBC network. When the recorded ON mBC is clamped at this potential, no current will flow through the gap junctions. This means that currents originating from photoreceptor inputs to the central cell will also be shared by the coupled lattice, and the amount of current escape will depend on *g*^gap^.

Fig. 10D shows that electrical coupling shifts the reversal potential (*V*_rev_) of the light-induced *IV* relation to more positive potentials, due to the fact that the neighboring cells are not clamped to the same extent as the central ON mBC. The slope of the light-induced *IV* relations becomes steeper as the strength of coupling augments. This happens because the total rod-driven *IV* relation measured in the central ON mBC is the sum of the *IV* relations of the central neuron and the relations originating from neighboring cells, which are smaller and slightly shifted in the voltage axis. However, in the absence of *I*_KV_, there is no rectification of the rod-driven current.

When K^+^ channels are placed at the tips of the dendrites of all cells in the coupled syncytium (Fig. 10E-G), the activation of these channels in both the central ON mBC and in the coupled network enlarges the voltage difference between the somatic compartments of the central cell and its neighbors. In this condition, the effects of coupling will become asymmetrical: at more depolarized potentials, the activation of *I*_KV_ in the neighboring cells will lead to an increase in the rectification of the rod-driven *IV* relation (Fig. 10F). Here, too, the slope of the total light-induced *IV* relations changes with the strength of coupling, because it results from the sum of many *IV* relations slightly displaced in the voltage axis. Inward currents increase more at negative potentials relative to those at positive potentials because the activation of dendritic K^+^ channels in both the central and neighboring cells increases space clamp problems at depolarized levels. These curves strongly resemble the *IV* relations of Fig. 6E; the whole-cell *IV*s and leak-subtracted *IV*s for the same simulations are presented in Fig. 10E and G, respectively, as well as the range of *IV*s recorded from real cells (grey areas represent mean ± SD of 28 records from 12 cells). Taken together, these simulations suggest that although electrical coupling between mixed-input ON mBCs does not induce rectification of light-driven conductances *per se*, it does increase the effects of dendritic K^+^ channels on rod-driven *IV* relations.

Because electrical coupling slows down cells due to the increase in capacitive load, we measured the response peak latency in the coupled network with and without dendritic K^+^ channels (Fig. 10H) to a 100 mV voltage step applied to the central ON mBC of the lattice (spread of potential for the same experiment is depicted in Fig. 10B). Similar to what happens in isolated cells (Fig. 9A and D), adding a voltage-gated conductance to the dendritic tips of ON mBCs (red traces) speeds up responses of the coupled cells considerably, and decreases the latency differences between the g_gap_ values used. Since the gap junctional conductance of these neurons changes by up to two fold with light adaptation (Arai et al., 2010), dendritic *I*_KV_ could provide a way of keeping responses of the ON mBC lattice equally fast, regardless of the adaptive state of the retina.

### The Resistance of Distal Dendritic Compartments Must Be High

For our model to reproduce the voltage-clamp experimental data (Fig. 6A-B and 9G-H), we needed to simulate a high resistance compartment between the soma and the dendritic tip. This was achieved by working with very fine secondary dendrites and shafts (0.01-μm in diameter) – larger dendrites failed to yield the same results (*not shown*). Although a narrowing of the secondary dendrites in relation to the dendritic tips was described in ON mBCs of the smooth dogfish (Witkovsky and Stell, 1973), goldfish (Ishida et al., 1980; Klooster et al., 2001; Stell, 1978) and rudd (Scholes, 1975), as well as in rod BCs of the guinea pig and rabbit retinas (Ladman, 1958; Sjostrand, 1998a; Sjostrand, 1998b), it is unlikely that dendritic diameters are as small as in our simulations. Rather, one needs to consider the influence of the extracellular resistance of the rod pedicle onto this system.

Fig. 11A shows a schematic drawing of the circuit involved in the rectification of the light-driven conductance of ON mBCs. In this drawing, secondary dendrites and dendritic shafts were lumped into one resistor. The resistance of the secondary dendrites and shafts at rod spherules (2) and cone pedicles (2’) is larger than that of the soma and primary dendrites (1). In a voltage-clamp experiment, current flowing from the soma (1) to the tips of the dendrites (3 and 3’) induces a voltage drop between the somatic compartment (as measured in the nodes *a* and *a’*) to the tips (measured in nodes *b* and *b’*). The residual voltage that reaches the tips opens the voltage-gated K^+^ channels located at the tips, and voltage control of the dendritic tips is lost.

**Figure 11.**
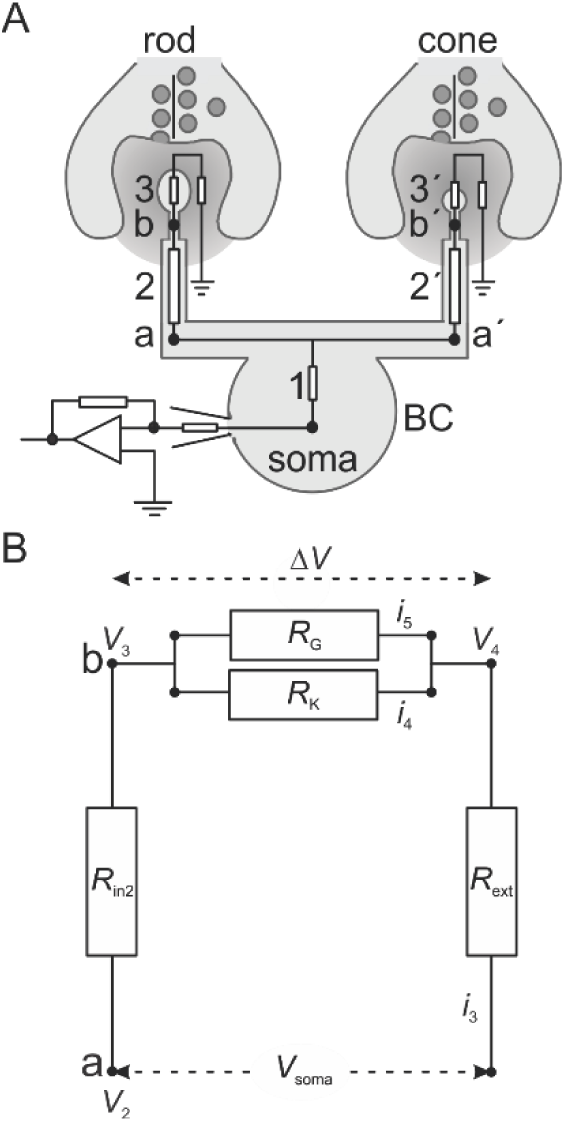
Electrical circuit involved in the rectification of light-induced *IV*s. **A)** Schematic drawing of the circuit involved in the rectification of the light-driven conductance of ON mBCs. **B)** Simplified equivalent scheme of an ON mBC. *V*_soma_, membrane potential at the soma; *V*_2_, potential at the secondary dendrites; *V*3, potential at the tips of the ON mBC dendrites; *R*_m_, membrane resistance of the soma and the primary dendrites; *R*_in2_, intracellular resistance of the dendrites invaginating rods; *R*_G_, TRPM1 conductance; *R*_K_, dendritic K^+^ conductance; *R*_ext_, resistance of the extracellular space; *i*_3_, current flowing through *R*_G_ and *R*_K_ at the dendritic tips. For formulas and further explanation, please see text.

Fig. 11B shows the simplified equivalent scheme of an ON mBC dendrite contacting a rod *in situ* (the left half of Fig. 11A). There are two variable resistors in this dendrite: (a) *R*_K_, representing the voltage-gated K^+^ conductance, and (b) *R*_G_, the glutamate-gated conductance through mGluR6/TRPM1. The total current flowing in the dendrite, *i*_1_, will flow through *R*_in2_, through the resistances at the dendritic tips *R*G and *R*_K_, and through the extracellular compartment (*R*_ext_). With this scheme, one can calculate whether *R*_in2_ is determinant for the membrane voltage over the TRPM1 channels (Δ*V*), or whether the extracellular resistance, *R*_ext_, can influence Δ*V*. In other words, one needs to compare this system when *R*_in2_ is high and *R*_ext_ is low and *vice-versa*.

The relationship between *V*_2_, the voltage at the secondary dendrites, Δ*V*, *R*_in2_ and *R*ext can be calculated by means of the following equations:

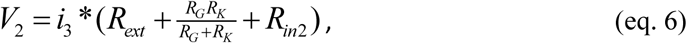

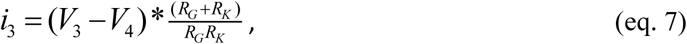

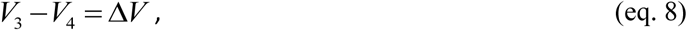

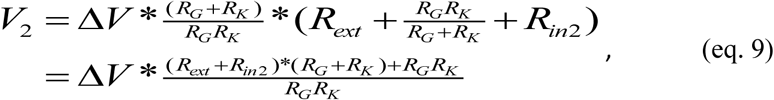

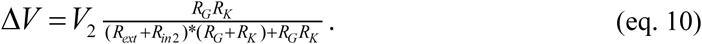

Eq. 10 shows that *R*_in2_ and *R*_ext_ are fully interchangeable: the space clamp problem will be similar when either the intracellular or extracellular resistance is high (or any combination in between). Therefore, even though ON mBCs might have relatively small dendritic diameters, there might also be a high extracellular resistance in the rod spherule, which would contribute to the results shown here. Such high resistance has been suggested to exist in the cone pedicle and to determine interactions between horizontal cells and cones in the outer retina (Kamermans et al., 2001; Klaassen et al., 2012; Vroman et al., 2013; Vroman et al., 2014). In the rod spherule, it creates an impedance mismatch between the sites of synaptic input and the cell soma, preventing proper space clamp at depolarized potentials.

## Discussion

Here we show that voltage-gated K^+^ channels, located at the dendritic tips of ON mBCs, act as a gain control mechanism for the rod-ON mBC synapse, by speeding up light responses of coupled cells and repolarizing their membrane potential in an intensity-dependent fashion. We will next discuss the consequences of this organization in visual processing.

### Bipolar Cells Are Not Isopotential

Most arguments about bipolar cells being isopotential come from experiments conducted on isolated neurons (Attwell et al., 1987; Lasater et al., 1984). However, the isolation procedure may lead to amputation of peripheral dendrites, since a high percentage of cells in these preparations that lack responses to glutamate (Lasater et al., 1984) and GABA (Kaneko et al., 1991). Even though anisopotentiality is usually thought of as a property of large CNS neurons, there are several examples in literature regarding compartmentalization of active conductances in bipolar cells. Some voltage-gated K^+^ channel subunits are preferentially expressed at the dendritic tips of murine rod bipolar neurons (Klumpp et al., 1995; Pinto and Klumpp, 1998), voltage-gated Na^+^ channels were identified in goldfish cone-driven bipolar cell dendrites and somas (Zenisek et al., 2001), and large Ca^2+^, Ca^2+^-dependent K^+^ and Cl^−^ currents originate in the giant synaptic terminals of goldfish ON mBCs (Burrone and Lagnado, 1997; Kaneko et al., 1991; Kaneko and Tachibana, 1985; Okada et al., 1995; Sakaba et al., 1997; Tachibana et al., 1993; Zenisek and Matthews, 1997) and in rat rod bipolar cells (Singer and Diamond, 2003).

In agreement with this idea, our simulations show that due to the nature of the synaptic inputs to ON mBCs (mGluR6 at the dendrites contacting rods and EAAT5 at the dendrites contacting cones), the dendritic tips contacting rods might be slightly more depolarized than the cell soma, while the dendrites contacting cones would be slightly more hyperpolarized than the soma (not shown). This inhomogeneity in a small neuron such as the goldfish ON mBC has important functional consequences, because it leads to a larger activation of dendritic K^+^ channels at the sites of rod synaptic input as compared to the sites of cone input. This way, dendritic *I*_KV_ will be more efficient in regulating the gain of the rod-ON mBC synapse than in regulating the gain of the cone-ON mBC synapse, having a larger influence in visual processing at scotopic and mesopic levels.

### Dendritic K^+^ Channels Are a Gain Control Mechanism

The rod-bipolar cell synapse has high gain in order to allow large threshold responses in the scotopic range (Ashmore and Falk, 1980; Capovilla et al., 1987; Copenhagen et al., 1990; Falk, 1988). Such an amplification at this synapse is necessary because rod voltage responses at threshold are extremely small (Copenhagen and Owen, 1976; Fain, 1975; Fain, 1976), slow, and buried in noise (Field et al., 2005; Taylor and Smith, 2004). Amplification of rod signals increases the responses of second-order neurons, but it also implies in amplification of noise levels (Field et al., 2005; Taylor and Smith, 2004). Such amplification is however not necessary for light intensities that generate sizeable responses from rods and that stimulate cones. The gain of the rod-bipolar cell synapse is therefore not static and changes with stimulus intensity and/or background light levels (Ashmore and Falk, 1980; Yang and Wu, 1997): the more photons available, the lower the gain. One of the mechanisms responsible for the high gain of this synapse and for eliminating some of its noise is the mGluR6 cascade itself (Falk, 1988; Sampath and Rieke, 2004; Shiells, 1994; Shiells and Falk, 1995). The mechanism responsible for the gain decrease at higher light levels is unknown – for this purpose, multiple mechanisms could be employed, such as feedback from horizontal cells onto rods (Thoreson et al., 2008) and intrinsic regulation of the mGluR6 cascade (Snellman et al 2008).

We propose that one such mechanisms is the activation of voltage-gated K^+^ currents in the vicinity of synaptic input sites. In the dark, the mGluR6 pathway is activated in a sustained manner, keeping the TRPM1 channels in the ON mBC membrane almost closed. Since in this condition the tips of the dendrites are relatively hyperpolarized, *I*_KV_ is hardly activated. This makes the resistance of the tips of the dendrites higher. In this condition, a small modulation of the mGluR6 pathway will lead to signal transmission to the soma, since the resistance of the tip is balanced to the resistance of the secondary dendrite. At higher light levels, the mGluR6 pathway will be less activated, and the tips of the dendrites will depolarize. This will in turn activate *I*_KV_, and the overall result will be a reduction of the resistance of the tips. Because in this condition the balance between the resistance of the tips and the resistance of the secondary dendrites is less optimal, the signals from rods to the ON mBC will be transmitted with lower gain. Interestingly, this system could also modify the gain of rod signals during light and dark adaptation, since voltage-gated K^+^ conductances in ON mBCs are enhanced by dopamine (Fan and Yazulla, 1999; Fan and Yazulla, 2001; Yazulla et al., 2001), and that dopamine levels are higher in the light-adapted retina (Djamgoz and Wagner, 1992; Witkovsky and Dearry, 1991).

This system has additional intriguing features. First, the interaction between dendritic K^+^ channels and the rod-driven conductance shown here not only speeds up repolarization of the ON mBC directly stimulated by light, but also contributes to signal spread within the network of coupled ON mBCs. The large convergence of rods onto ON mBCs and the fact that the latter are electrically coupled make them poor candidates for single-photon detectors, unlike mammalian rod-driven bipolar cells, since (i) photoreceptor convergence can potentially lead to a decrease in the influence of each single receptor onto the membrane potential of the second-order neuron (Falk, 1988), and (ii) electrical coupling decreases the flash sensitivity of individual cells due to current escape via the gap junctions (Hornstein et al., 2005). Rather, this combination of large receptive fields and transient responses would tune ON mBCs for the detection of low spatial frequencies and middle to high temporal frequencies, which could in turn be beneficial for downstream neurons involved in motion detection.

Finally, although mammalian rod-driven bipolar cells are not electrically coupled, they share with ON mBCs the same machinery for the generation and gain control of the light response. In this respect, the results shown here are relevant for both mammalian and non-mammalian retinas. Because K^+^ currents do not affect small (2-5 mV) voltage responses, at very low light levels, ON mBCs are allowed to integrate photons over larger periods. As more and more photons become available, the proposed mechanism will quickly change the gain of the synapse locally to prevent saturation, while allowing signals from other rods to activate the ON mBC with the highest possible gain. Since the activation of these K^+^ channels allows the membrane voltage to respond quicker to the changes in current flow (Mao et al., 1998; Mao et al., 2002), a consequence of this is that individual dendritic responses become shorter than the rod response itself, eliminating temporal redundancy in the neural code already at the first synapse. In fact, rod-driven ON BC responses are much faster than rod light responses in both lower vertebrates and mammals (Field et al., 2005).

## Author contributions

C.J. and M.K. designed the experiments, C.J. and J.K. performed the experiments, C.J. and M.K. performed the simulations, C.J., J.K. and M.K. wrote the manuscript. All authors approved the final version of the manuscript.

## Acknowledgements

This work was supported by Conselho Nacional de Desenvolvimento Científico e Tecnológico (CNPq, 200915/98-3), Fundação de Amparo à Pesquisa do Estado de São Paulo, Brazil (FAPESP, 2010/16469-0), and The European Office of Aerospace Research & Development (FA8655-05-C-4018). The authors would like to thank Drs. Leon Lagnado, Robert G. Smith and Henrique von Gersdorff for helpful discussions.

## Abbreviations List

ACPT-1: (1S,3R,4S)-1-Aminocyclopentane-1,3,4-tricarboxylic acid
BC: bipolar cell
*C*_m_: specific membrane capacitance
DAB: diaminobenzidine
DL-AP4: DL-2-Amino-4-phosphonobutyric acid
EAAT5: excitatory amino acid (glutamate) transporter 5
*E*_cone_: reversal potential of the cone-driven conductance
*E*_K_: reversal potential of the potassium conductance
*E*_leak_: reversal potential of the leak conductance
*E*_rod_: reversal potential of the rod-driven conductance
EM: electron microscopy
FWHM: full width at half-maximum
*g*_gap_: gap-junctional conductance
HC: horizontal cell
*I*_gap_: gap-junctional current
*I*_A_: transient voltage-gated potassium current
*I*_KV_: voltage-gated potassium current
INL: inner nuclear layer
*IV*: current-voltage relationship
LM: light microscopy
LY: lucifer yellow
mGluR: metabotropic glutamate receptor
mGluR6: metabotropic glutamate receptor 6
NGS: normal goat serum
ONL: outer nuclear layer
ON BC: depolarizing bipolar cell
ON mBC: depolarizing mixed-input bipolar cell
OPL: outer plexiform layer
PB: phosphate buffer
PKC: phosphokinase C
PTX: picrotoxin
*R*_a_: specific cytoplasmic resistivity
*R*_in_: input resistance
*R*_m_: specific membrane resistance
STRY: strychnine
*V*hold: holding membrane potential
*V*_rest_: resting membrane potential
*V*_soma_: somatic membrane potential
*V*_step_: voltage step
*V*_tip_: membrane potential at the dendritic tips
TRPM1: transient receptor potential channel, type melastatin-1

